# TRIM37 employs peptide motif recognition and substrate-dependent oligomerization to prevent ectopic spindle pole assembly

**DOI:** 10.1101/2024.10.09.617493

**Authors:** Andrew Bellaart, Amanda Brambila, Jiawei Xu, Francisco Mendez Diaz, Amar Deep, John Anzola, Franz Meitinger, Midori Ohta, Kevin D. Corbett, Arshad Desai, Karen Oegema

## Abstract

Tightly controlled duplication of centrosomes, the major microtubule-organizing centers of animal cells, ensures bipolarity of the mitotic spindle and accurate chromosome segregation. The RBCC (RING-B-box-coiled coil) ubiquitin ligase TRIM37, whose loss is associated with elevated chromosome missegregation and the tumor-prone developmental human disorder Mulibrey nanism, prevents the formation of ectopic spindle poles that assemble around structured condensates containing the centrosomal protein centrobin. Here, we show that TRIM37’s TRAF domain, unique in the extended TRIM family, engages peptide motifs in centrobin to suppress condensate formation. TRIM proteins form anti-parallel coiled-coil dimers with RING–B-box domains on each end. Oligomerization due to RING–RING interactions and conformational regulation by B-box-2–B-box-2 interfaces are critical for TRIM37 to suppress centrobin condensate formation. These results indicate that, analogous to anti-viral TRIM ligases, TRIM37 activation is linked to the detection of oligomerized substrates. Thus, TRIM37 couples peptide motif recognition and substrate-dependent oligomerization to effect ubiquitination-mediated clearance of ectopic centrosomal protein assemblies.

## MAIN

Centrosomes are the major microtubule-organizing centers of animal cells and comprise a centriolar core that recruits a pericentriolar material matrix (PCM matrix) to nucleate and anchor microtubules. Centrosome duplication is tightly controlled so that mitotic cells have exactly two centrosomes that catalyze microtubule assembly to form the two poles of the bipolar spindle (Banterle and Gonczy, 2017; Gomes Pereira et al., 2021). Centrosomes are multi-gigadalton macromolecular assemblies comprising ∼150 proteins organized into multiple substructures (Laporte et al., 2024; LeGuennec et al., 2021; Ma et al., 2023). Given the complex interconnected nature of the centrosomal protein network, a key question is how rogue collections of centrosomal proteins are prevented from assembling in the cytoplasm to form ectopic structures with microtubule-organizing capacity.

Recent work identified the ubiquitin ligase TRIM37, a member of the <u>TRI</u>partite <u>M</u>otif protein family, as a key player in defending against the formation of ectopic assemblies of centrosomal proteins (Balestra et al., 2021; Balestra et al., 2013; Meitinger et al., 2016; Meitinger et al., 2021; Meitinger et al., 2020; Yeow et al., 2020). TRIM family ligases are defined by an RBCC domain composed of a RING domain, B-box, and an anti-parallel coiled-coil (Esposito et al., 2017; Fiorentini et al., 2020; Koepke et al., 2021). Unique among TRIM proteins, TRIM37 also harbors a TRAF domain, first identified in TNF receptor-associated factors, that is predicted to bind to peptide ligands (Park, 2021; Zapata et al., 2007). In cells with centrioles, TRIM37 loss leads to the formation of a single large and highly ordered assembly (termed a “condensate”) containing the centrosomal protein centrobin (encoded by *CNTROB*), as well as an array of smaller foci containing the centriolar protein centrin. In ∼25% of mitotic cells, the centrobin condensate forms an ectopic spindle pole, leading to transient or persistent multipolarity and elevated chromosome missegregation (Balestra et al., 2021; Balestra et al., 2013; Meitinger et al., 2021). *CNTROB* deletion suppresses ectopic spindle pole formation in *TRIM37Δ* cells (Meitinger et al., 2021). Thus, ectopic centrobin-scaffolded condensates function as aneuploidy generators and may underlie the high prevalence of tumors in patients with the TRIM37 loss-of-function disorder Mulibrey nanism (Avela et al., 2000; Karlberg et al., 2009).

A critical knowledge gap we address is how TRIM37 recognizes nascent centrobin assemblies and promotes their disassembly to suppress condensate formation. We identify short peptide motifs in a predicted disordered C-terminal region of centrobin that engage TRIM37’s TRAF domain and are essential for TRIM37 to bind centrobin and prevent its ectopic assembly *in vivo*. In addition to TRAF domain-mediated motif recognition, TRIM37 detects substrate oligomerization to mediate clearance. TRIM37 is predicted to form anti-parallel coiled-coil dimers that place their RING-B-box-2 domains on opposing ends (Esposito et al., 2017; Sanchez et al., 2014). As RING domain dimerization is typically required for ubiquitin ligase activation (Fiorentini et al., 2020), the RING-B-box-2 region must further oligomerize, potentially facilitated by binding to oligomeric substrates, for efficient substrate ubiquitination. Using engineered recombinant miniTRIM37 constructs, we show that the RING domain, but not the B-box-2, represents the primary TRIM37 oligomerization interface. The B-box-2 interface controls the conformation of TRIM37 oligomers. *In vivo*, the RING and B-box interfaces are both essential to recognize centrobin assemblies and prevent condensate formation. Collectively, these results indicate that, analogous to the anti-viral TRIM ligase TRIM5α (Spada et al., 2024), TRIM37 employs a combination of substrate motif binding and substrate-dependent oligomerization to effect ubiquitination-mediated clearance of ectopic centrosomal protein assemblies.

## RESULTS

### TRIM37 can act in the cytosol to prevent the formation of centrobin condensates

In fibroblasts isolated from Mulibrey nanism patients and *TRIM37Δ* RPE1 cells, which both lack TRIM37, most cells possess a single highly-ordered non-centrosomal condensate containing the centriolar protein centrobin. In ∼25% of mitotic cells, the centrobin-containing condensate acquires the ability to nucleate microtubules and forms an ectopic spindle pole, leading to transient or persistent multipolarity and elevated chromosome missegregation (**Fig. 1A**; (Balestra et al., 2021; Meitinger et al., 2021)). Consistent with an essential scaffolding role for centrobin in condensate formation, knockout of the gene encoding centrobin (*CNTROB*) in *TRIM37Δ* cells eliminated detectable centrobin signal and ectopic spindle poles (Balestra et al., 2021; Meitinger et al., 2021). While prior immunofluorescence suggested that PLK4 may also localize to centrobin condensates (Meitinger 2021, Meitinger 2020), analysis of an inducible *PLK4* knockout in a *TRIM37Δ*; *p53sh* cell line suggested that the detected PLK4 was due to cross-reactivity with another protein in the condensate (**Fig. S1A-C**). We therefore conclude that the centrobin-scaffolded condensates do not contain PLK4.

**Figure 1.**
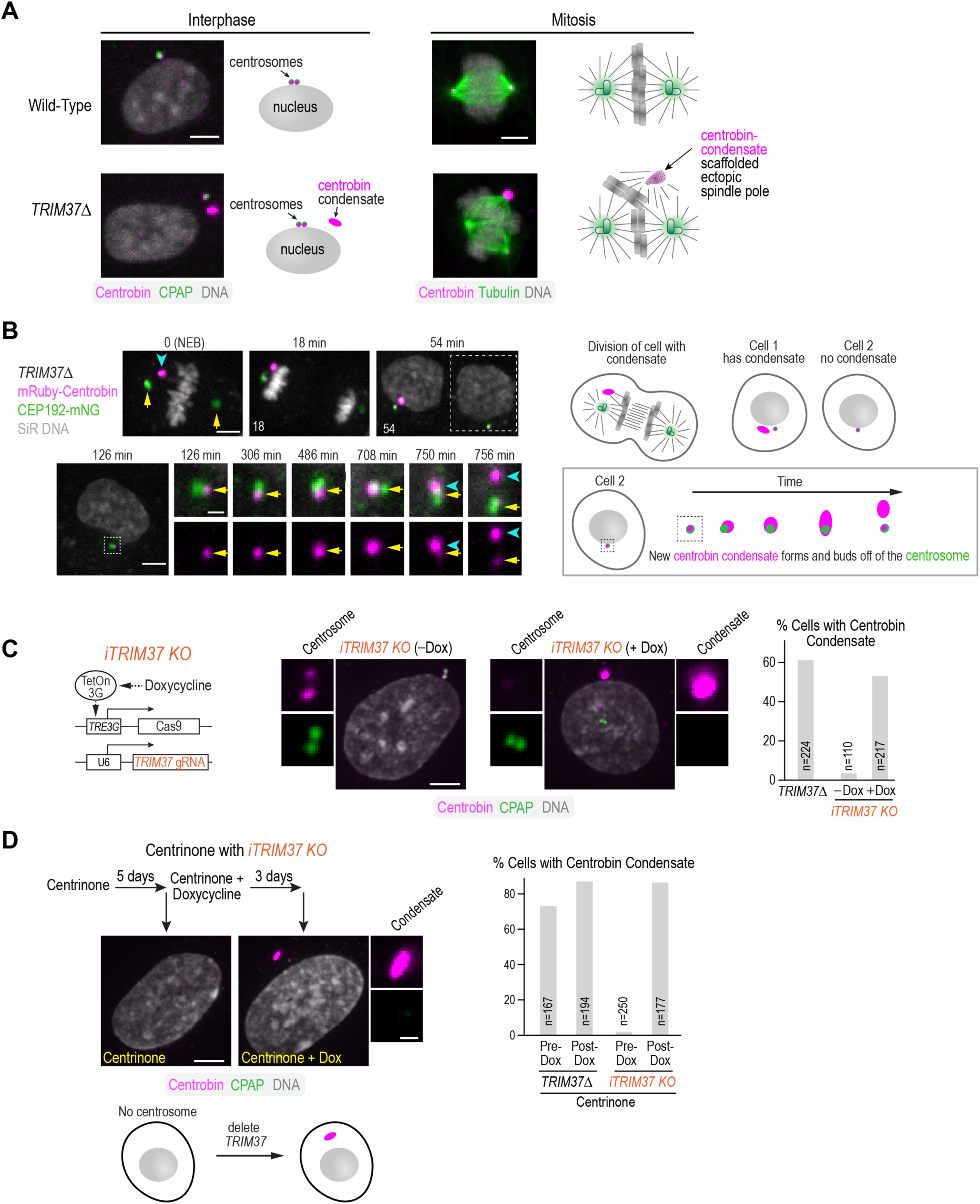
TRIM37 prevents the formation of centrobin condensates in the cytosol independently of centrosomes. **(A)** Representative images and schematic summary of phenotypes in interphase and mitotic wildtype and *TRIM37Δ* RPE1 cells. Loss of TRIM37 leads to the formation of a centrobin-containing condensate. Upon mitotic entry, centrobin condensates form ectopic spindle poles at an appreciable frequency, leading to elevated rates of chromosome missegregation. **(B)** Imaging of a *TRIM37Δ* RPE1 cell expressing mNG-tagged CEP192 to mark the centrosomes and mRuby-tagged centrobin; SiR-DNA was added to visualize chromosomes—schematics on the right aid in the interpretation of the images. Since *TRIM37Δ* cells typically have a single condensate, following mitosis, the daughter cell on the left inherits the condensate, while the one on the right *(boxed with a dashed line in the 54 min panel*) does not. The cell that did not inherit the condensate was imaged live (*row below*); the region containing the centrosome is magnified on the right; top row is a merge of the CEP192 centrosome marker and centrobin; bottom row shows only centrobin. Centrosomes, visualized by CEP192, are indicated with yellow arrows and the newly forming centrobin condensate with cyan arrowheads. Times are relative to metaphase in the mother cell. A similar phenomenon was observed in 4 cells. **(C)** (*left*) Schematic summary of the approach used to inducibly knockout *TRIM37* (see also **Fig. S1D,E**); (*middle*) immunofluorescence images of an inducible TRIM37 knockout (*iTRIM37KO*) cell line without and with doxycycline-induced, Cas9-mediated knockout. Cells were stained for centrobin and the centriolar marker CPAP. Centrosomes are magnified to the left of the lower magnification view and, when present, centrobin condensates to the right. (*right*) Graph plotting the frequency of centrobin condensates for the indicated conditions. *n* is the number of cells analyzed. **(D)** (*top left*) Experimental scheme for analyzing the effect of knocking out *TRIM37* in cells lacking centrosomes, following PLK4 inhibition with centrinone, and images of cells treated as indicated and labeled for centrobin and CPAP; absence of focal CPAP staining indicates absence of centrioles (see also **Fig. S1F**). A magnified view of the condensate shows centrobin staining on top (*magenta*) and CPAP signal, which is absent, on the bottom. (*right*) Graph comparing the frequency of condensate formation for the indicated conditions. *n* is number of cells analyzed. Scale bars are 5µm in panels showing lower magnification views and 1 µm for centrosome and condensate blowups.

Centrobin condensates form when *TRIM37* is deleted. Complementing *TRIM37Δ* cells with epitope-tagged wild-type TRIM37 prevents condensate formation, whereas complementation with ligase-mutant (C18R) TRIM37 does not (Meitinger et al., 2021; Meitinger et al., 2020), indicating that the ubiquitin ligase activity of TRIM37 is essential for suppressing condensate formation. Although it cannot suppress condensate formation, fluorescently tagged ligase-mutant TRIM37 (Lig^mut^ TRIM37-mNG) stably binds to condensates and allows monitoring of their dynamics (Meitinger et al., 2021; Meitinger et al., 2020). When a cell with a single condensate divides, one cell inherits the condensate, and the other does not (**Fig. 1B**). In cells born without a condensate, Lig^mut^ TRIM37-mNG was observed to hyper-accumulate around the centrosome and then bud off, suggesting that new condensates form by budding off of centrosomes (Meitinger et al., 2021). To confirm that this is also true in the absence of the mutant ligase, we imaged mRuby-centrobin in cells expressing CEP192-mNG to mark the centrosomes. In *TRIM37Δ* daughter cells that failed to inherit a condensate, mRuby-centrobin hyper-accumulated around the centrosome and then budded off to form a condensate (**Fig. 1B**). This observation raises the question of why there is only a single condensate, and whether new condensates can spontaneously form in the cytoplasm or if they must be ‘nucleated’ by an existing structure like the centrosome or an existing condensate.

To determine whether cells lacking a centriole or condensate can spontaneously form a condensate, we engineered and validated an inducible *TRIM37* knockout (**Fig. 1C; Fig. S1D,E**). We first depleted centrosomes from cells by treating them with the PLK4 inhibitor centrinone (**Fig. S1F**; (Wong et al., 2015)) and then added doxycycline to induce the *TRIM37* knockout (**Fig. 1D**). If centrosomes nucleate condensate formation, we would expect that either a condensate would be unable to form, or multiple condensates would form and grow simultaneously. In contrast to this expectation, inducibly deleting *TRIM37* after centrosome loss resulted in the formation of centrobin condensates at a frequency similar to that in cells constitutively lacking *TRIM37* with centrosomes (**Fig. 1D**). These results suggest that loss of *TRIM37* initiates the assembly of ordered centrobin-containing condensates. Although condensates preferentially form at centrosomes when they are present, possibly building on a scaffold that exists there, a condensate can also spontaneously form in the cytoplasm. The fact that only one condensate typically forms, even in cells that lack a centrosome, suggests that condensates are nucleated infrequently in cells that do not have one and that it is easier to add to an existing condensate than to initiate a new one. It also indicates that TRIM37 can suppress the formation of oligomeric centrobin-containing assemblies both at centrosomes and in the cytoplasm.

### The TRIM37 TRAF domain is required for centrobin binding and condensate clearance

TRIM family ligases are defined by an RBCC domain containing a RING domain, B-box, and an anti-parallel coiled-coil (Esposito et al., 2017; Fiorentini et al., 2020). AlphaFold modeling of TRIM37 suggested that, as in other TRIM proteins, the TRIM37 coiled-coil forms a T-shaped antiparallel dimer that places its two RING-B-box-2 domains on opposite sides (**Fig. 2A; Fig. S2A,B**). Unique among TRIM family proteins, TRIM37 also harbors a TRAF domain (**Fig. 2A**; **Fig. S2C**). TRAF domains, first identified in TNF receptor-associated factors, bind short peptide sequences, with different TRAF domains exhibiting binding specificity for different peptide motifs (Park, 2021; Zapata et al., 2007). AlphaFold predicts that the two TRAF domains are positioned on the stem of the T, just below the extended antiparallel coiled-coil (**Fig. 2A**). We hypothesized that the TRIM37 TRAF domain recognizes specific peptide motifs in targets such as centrobin to position them for ubiquitination by the RING-E2 complex. To test this model, we designed a TRAF domain mutant based on structural homology to USP7, which binds peptide ligands in p53 (Hu et al., 2006; Sheng et al., 2006). Specifically, we mutated a critical exposed tryptophan residue in the TRIM37 TRAF domain to alanine (**Fig. 2B**; W373>A; referred to as TRAF^mut^; (Meitinger et al., 2020)), which is predicted to disrupt its ability to engage its peptide ligands. After confirming the expression of transgene-encoded WT, Lig^mut^, and TRAF^mut^ TRIM37 proteins in *TRIM37Δ* cells (**Fig. 2B**), we analyzed the frequency of centrobin condensate formation. In contrast to the WT transgene, whose expression completely suppressed centrobin condensate formation, expressing TRAF^mut^ or Lig^mut^ TRIM37 did not alter the frequency of condensate formation compared to the *TRIM37Δ* background (**Fig. 2C-E**). Thus, the TRAF domain interface predicted to engage peptide ligands is essential for TRIM37 to suppress the formation of centrobin condensates.

**Figure 2.**
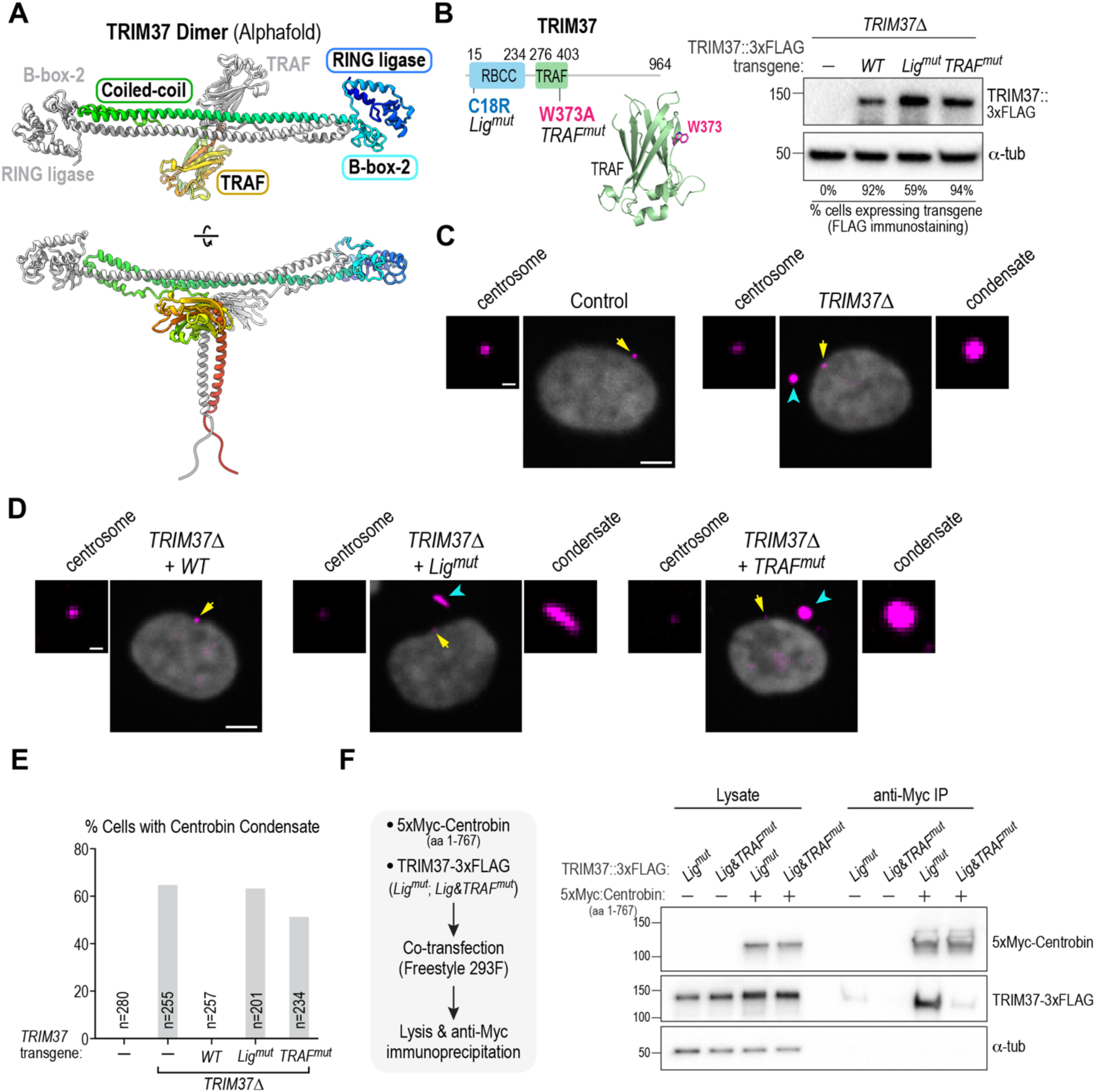
The TRAF domain of TRIM37 is required to prevent centrobin condensate formation *in vivo* and for binding of TRIM37 to centrobin. **(A)** AlphaFold model of the anti-parallel TRIM37 dimer. The RING and B-box-2 domains are on opposite ends of the dimer, and the TRAF domains are positioned just below the mid-point of the anti-parallel coiled-coil. One TRIM37 monomer is colored from the N-terminus in blue to the C-terminus in red; the second TRIM37 monomer is colored grey. See also **Fig. S2A,B**. **(B)** (*left*) Schematic of TRIM37 along with an AlphaFold model of the TRAF domain highlighting the tryptophan residue (W373) that was mutated to prevent putative peptide ligand engagement (see also **Fig. S2C**). (*right*) Immunoblot of *TRIM37Δ* RPE1 cells engineered to express the indicated TRIM37 variants using lentiviral transduction. The TRIM37 transgenes include a FLAG tag, which is immunoblotted; α-tubulin serves as a loading control. Numbers below the lanes indicate the percentage of cells in the transduced pool that express the transgene, as assessed by anti-FLAG immunostaining. **(C) & (D)** Images of control and *TRIM37Δ* RPE1 cells (*C*) and of *TRIM37Δ* cells expressing the indicated *TRIM37* transgenes (*D*). Cells were labeled for centrobin and DNA; centrosomes (*yellow arrows*) and condensates (*cyan arrowhead*s) are indicated on the images. Centrosomes are magnified to the left of the lower magnification view and, when present, centrobin condensates to the right. **(E)** Frequency of centrobin condensate formation for the indicated conditions. *n* is the number of cells analyzed. **(F)** (*left*) Experimental schematic of analysis of TRIM37 interaction with centrobin following co-expression in FreeStyle 293F cells. The ligase activity of TRIM37 was mutated to enable robust expression and assessment of binding. The centrobin fragment (1-767) is soluble and contains the TRIM37-binding region (see Fig. 3A & **Fig 4A**), facilitating the binding analysis. (*right*) Immunoblot of Centrobin (1-767), detected using the Myc epitope tag, and TRIM37, detected using the FLAG epitope tag, in cell lysates and following anti-Myc immunoprecipitation. α-tubulin serves as a loading control for the input lysates. Scale bars in panels *C* and *D*, 5 µm (*lower magnification views*) and 1 µm (*centrosome and condensate blowups*).

To address whether the TRIM37 TRAF domain is essential for TRIM37 to bind to centrobin, we employed a human cell co-expression approach (**Fig. 2F**; (Meitinger et al., 2021)). Since full-length centrobin is largely insoluble, we assessed binding to a centrobin truncation (aa 1-767) that removes 125 aa from its C-terminus. Since TRIM37 negatively autoregulates its own abundance (Meitinger et al., 2020), we employed Lig^mut^ TRIM37 (C18R), which is present at higher levels than WT TRIM37, to analyze centrobin binding. Whereas Lig^mut^ TRIM37(C18R) bound robustly to centrobin 1-767, the binding of Lig & TRAF^mut^ TRIM37(C18R; W373A) was significantly reduced (**Fig. 2F**). We conclude that the TRAF domain-peptide ligand interface is necessary for TRIM37 to bind to centrobin and prevent the formation of centrobin condensates.

### The TRIM37 TRAF domain recognizes specific peptide motifs in centrobin

To delineate the molecular basis for the recognition of centrobin by the TRAF domain, we used a co-expression-based binding assay to narrow down the region of centrobin required for association with TRIM37 (**Fig. 3A**). This effort revealed that the TRIM37 binding region is located between aa 576 & 767 of centrobin. AlphaFold modeling predicted three potential peptide binding motifs for the TRAF domain within this region (**Fig. 3B**). Motif 1 was predicted with the highest confidence and motif 3 with the lowest confidence; two other predicted binding motifs (motifs 2.1 and 2.2) are overlapping (**Fig. 3B; Fig. S3A**). To determine if these peptide motifs are important for TRAF domain-mediated recognition of centrobin, we engineered a mutant centrobin altering 18 predicted interface residues in all motifs (referred to as TBM^mut^, for <u>T</u>RIM37 <u>b</u>inding <u>m</u>otif mutant) (**Fig. 3B,C**). Mutation of the three identified motifs in centrobin largely eliminated TRIM37 binding, highlighting their importance in the recognition of centrobin by TRIM37 (**Fig. 3C**). To directly assess motif binding, we performed fluorescence polarization-based peptide binding assays with purified recombinant WT or mutant (W373>A) TRIM37 TRAF domains (**Fig. 3D; Fig. S3B**). This analysis confirmed specific binding of motif 1 (K_d_ 14 µM) and weaker specific binding of motifs 2.1 and 2.2 (K_d_ 187 and 147 µM, respectively). By contrast, no specific binding was observed for motif 3. The relatively low affinities observed for these TRAF domain-peptide ligand motif interactions are in the range of those reported for other TRAF-peptide ligands (Hu et al., 2006; Sheng et al., 2006). Given the proximity of the two TRAF domains within the anti-parallel TRIM37 dimer (**Fig. 2A**), avidity from recognizing multiple low-affinity motifs may contribute to TRIM37 TRAF domain-mediated centrobin recognition.

**Figure 3.**
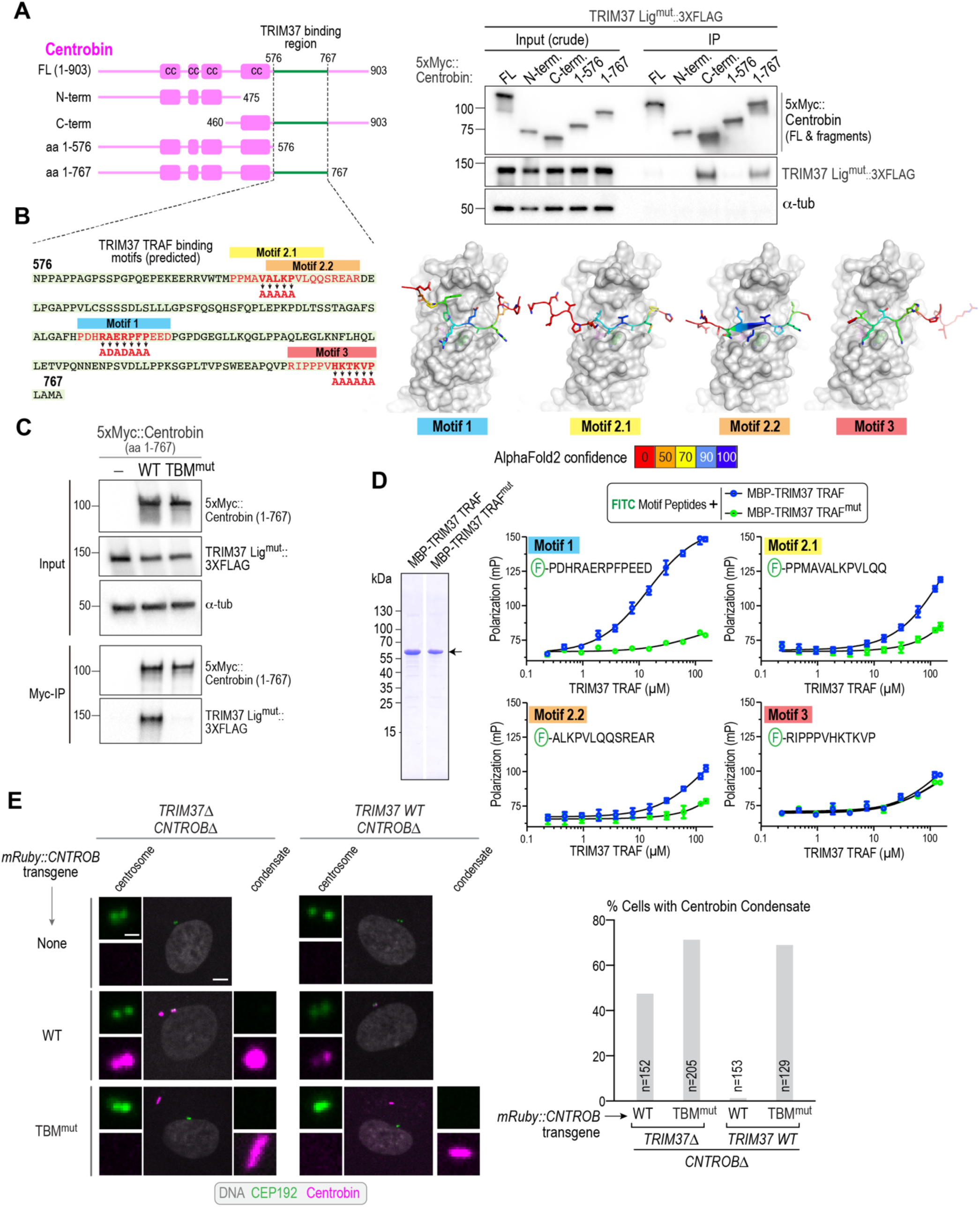
The TRIM37 TRAF domain recognizes specific peptide motifs in centrobin. **(A)** (*left*) Schematic of full-length (FL) and engineered centrobin fragments; *cc* refers to predicted coiled-coils. (*right*) Immunoblot analyzing binding of TRIM37 to centrobin fragments, performed as in Fig. 2F. Note that the input shown here is the crude extract, prior to centrifugation. For FL centrobin, which is largely insoluble, the clarified supernatant used for the anti-Myc immunoprecipitation is depleted of the TRIM37 ligase-mutant, which pellets with the insoluble centrobin assemblies; this explains the absence of a TRIM37 band in the FL centrobin immunoprecipitation. Similar results were observed in two independent experiments. **(B)** (*left*) Sequence of the TRIM37-binding region of centrobin, highlighting potential TRAF domain-binding motifs and mutations engineered to disrupt them; (*right*) AlphaFold models of centrobin motifs interfacing with the TRIM37 TRAF domain. The TRAF domain is shown in a space-filling view in gray; the motifs are colored by the AlphaFold prediction confidence score (see also **Fig. S3A**). Similar results were observed in two independent experiments. **(C)** Analysis of TRIM37 binding to wildtype or a mutant form of centrobin (1-767) in which the putative TRAF-binding motifs were mutated as indicated in Fig. 3B. The binding assay was conducted as in Fig. 2F. **(D)** (*left*) Coomassie-stained gel showing purified recombinant WT versus W373A-mutant TRAF domains (*arrow*). Both lanes shown are from the same gel; an intervening lane was removed (*white line*). (*right*) Analysis of binding of the indicated fluorescent peptides to the purified TRAF domains, monitored using fluorescence polarization (see also **Fig. S3B**). **(E)** (*left*) *In vivo* comparison of wildtype or TRIM37 binding motif-mutant centrobin expressed in either *TRIM37Δ;CNTROBΔ* or *TRIM37 WT; CNTROBΔ* cells (see also **Fig. S3C-E**). The centrobin transgenes included an mRuby tag for visualization. (*right*) Quantification of the frequency of centrobin condensate formation for the indicated conditions. *n* is the number of cells analyzed. Scale bars are 5µm in panels showing lower magnification views and 1 µm for centrosome and condensate blowups.

### TRAF domain-binding motifs in centrobin are required for TRIM37 to suppress centrobin condensate formation

If the TRAF binding motifs are important for TRIM37 to recognize centrobin *in vivo*, then their mutation would prevent TRIM37 from targeting centrobin and lead to the formation of centrobin condensates even when WT TRIM37 is present. To test this prediction, we employed CRISPR/Cas9 to generate stable *CNTROBΔ* and *CNTROBΔ;TRIM37Δ* RPE1 cell lines (**Fig. 3E**; **Fig. S3C-E**). Transgenes expressing a fusion of mRuby with WT centrobin, or a centrobin with its TRAF binding motifs mutated (TBM^mut^), were then introduced into the two cell lines by lentiviral transduction (**Fig. 3E**). In *TRIM37Δ* cells, both WT and TBM^mut^ centrobin formed condensates (**Fig. 3E**), indicating that the TRAF binding motif mutations do not prevent condensate formation. By contrast, only TBM^mut^ centrobin formed condensates in the presence of WT TRIM37 (**Fig. 3E**). These results indicate that TRAF domain-mediated binding of TRIM37 to specific motifs in centrobin is a critical step in preventing centrobin condensate formation. We additionally mutated only motif 1, which exhibited the most robust specific binding in the purified TRAF domain-peptide binding assay (**Fig. S4A,B**). While significantly higher than WT centrobin, the motif 1 centrobin mutant exhibited a much lower frequency of condensates than TBM^mut^ centrobin (9% vs 69%; **Fig. S4A,B**). This data suggests that multiple binding motifs contribute to centrobin recognition by TRIM37 dimers.

Collectively, these results highlight the importance of the TRIM37 TRAF domain in preventing the formation of centrobin condensates and delineate the molecular interfaces by which the TRAF domain recognizes centrobin.

### Centrobin oligomerization is important for its ubiquitination by TRIM37

TRIM37 suppresses the formation of centrobin condensates that function as non-centrosomal spindle poles (Balestra et al., 2021; Meitinger et al., 2021). The ability of TRIM37 to suppress the formation of large ectopic centrobin assemblies, while allowing centrobin to function at centrioles and in ciliogenesis (Karasu et al., 2022; Ogungbenro et al., 2018), suggests that TRIM37 might recognize and ubiquitinate large oligomers but not unassembled centrobin dimers in the cytosol. To determine if ubiquitination by TRIM37 of centrobin is linked to its oligomerization status, we first assessed the requirements for centrobin oligomerization using centrifugation. Our prior work showed that full-length centrobin expressed in human cells oligomerizes readily and fractionates into the pellet following centrifugation (Meitinger et al., 2021). In addition, when TRIM37 Lig^mut^ is co-expressed with full-length centrobin, TRIM37 Lig^mut^ co-sediments with the centrobin assemblies, mimicking the stable association of TRIM37 Lig^mut^ with centrobin-scaffolded condensates observed *in vivo* (Meitinger et al., 2021). To address requirements for centrobin oligomerization, we analyzed the fractionation of full-length centrobin and centrobin fragments into the supernatant and pellet fractions following centrifugation of cell extracts (**Fig. 4A**). While full-length centrobin largely fractionated into the pellet, all of the other centrobin fragments were soluble (**Fig. 4A**). We also used lentivirus-mediated transgene delivery to express the 1-767 fragment of centrobin *in vivo* and found that it neither localized to centrosomes nor formed condensates (*not shown*). These data indicate that centrobin oligomerization requires both N-terminal and C-terminal regions that are distinct from the TRIM37-binding region.

**Figure 4.**
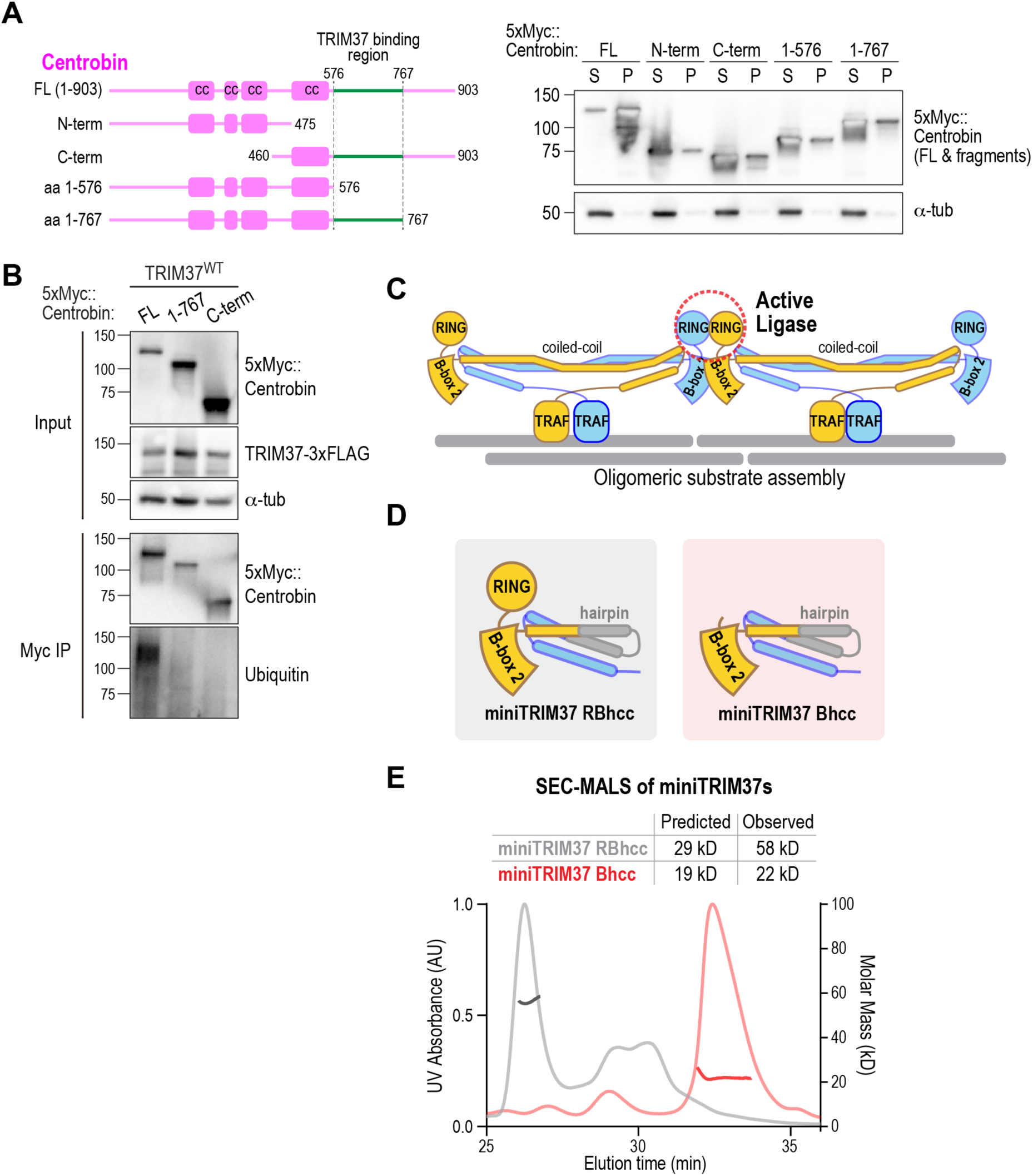
Centrobin oligomers are targeted for ubiquitination by TRIM37, whose oligomerization requires the RING domain. **(A)** (*left*) Schematic of full-length centrobin and fragments highlighting the TRIM37 binding region (*green*); (*right*) analysis of solubility of indicated centrobin variants expressed in Freestyle 293F cells. α-tubulin serves as a loading and solubility control. The blot was overexposed to highlight the signals in the supernatant (FL) and pellets (all other variants). **(B)** Analysis of centrobin fragment ubiquitination by WT TRIM37 following co-expression in FreeStyle 293F cells followed by immunoprecipitation and immunoblotting. Endogenous ubiquitin was detected in the immunoprecipitates using an anti-ubiquitin antibody. **(C)** Schematic model of substrate oligomerization-directed activation of TRIM37 ligase activity. **(D)** Schematics of mini-TRIM37 variants engineered to examine the roles of the RING and B-box-2 domains in oligomerization (see also *Fig. S4C*). **(E)** SEC-MALS analysis of purified recombinant miniTRIM37s. The numbers above indicate the predicted molecular weights from the primary sequences and the native molecular weights of the entities in the major UV peaks measured by SEC-MALS.

Next, we co-expressed WT TRIM37 with either full-length centrobin or N- or C-terminal truncations containing the TRIM37-binding region and analyzed their ubiquitination. While the small pool of full-length centrobin in the supernatant was robustly ubiquitinated by WT TRIM37, the two soluble fragments were not (**Fig. 4B**). As these two fragments both bind to TRIM37 and together span the entire centrobin sequence, these data suggest that centrobin oligomerization is important for its ubiquitination by TRIM37.

### The TRIM37 RING–B-box-2 oligomerizes primarily via a RING-RING interface

Next, we focused on addressing how oligomeric centrobin assemblies activate TRIM37-mediated ubiquitination. Prior work on TRIM proteins (Esposito et al., 2017; Fiorentini et al., 2020) supported by AlphaFold modeling (**Fig. 2A**) suggests that TRIM37 forms an antiparallel dimer with its two RING-B-box-2 domains positioned on opposite ends of a T-shaped dimer. As RING domain dimerization is important for activating the bound E2-ubiquitin complex for substrate ubiquitination (Fiorentini et al., 2020), further oligomerization of TRIM37 dimers fostered by binding to an oligomeric substrate could contribute to ligase activation and substrate ubiquitination (**Fig. 4C**). Such an activation mechanism has been elucidated for TRIM5α, which detects viral nucleocapsid shells in the cytosol and targets them for ubiquitin-mediated degradation (Ganser-Pornillos and Pornillos, 2019; Spada et al., 2024). Similar to TRIM37, TRIM5α has a RING–B-box-2-coiled coil (RBCC) domain that is followed, rather than by a TRAF domain, by a SPRY domain that recognizes the viral nucleocapsid. Capsid-driven oligomerization of TRIM5α is important for its activation. The ability of TRIM5α dimers to oligomerize in a manner that matches the geometry of the viral nucleocapsid requires an interaction interface on its B-box-2 domains, with a well-characterized point mutation in this interface disrupting oligomerization and the ability to restrict HIV infection (Ganser-Pornillos et al., 2011; Li and Sodroski, 2008; Wagner et al., 2016).

For TRIM5α, RING and B-box-2-mediated oligomerization was analyzed by engineering “miniTRIM” constructs that contained either a single RING–B-box-2 or B-box-2 domain only fused to a short hairpin coiled-coil from *T. thermophilus* seryl-tRNA synthetase (PDB ID 1SER, (Biou et al., 1994); (Wagner et al., 2016)). This approach allowed analysis of the RING and B-box-2 interfaces that mediate oligomerization independently of the extended anti-parallel coiled-coil of the native dimer. Both miniTRIM5α constructs were dimers, and a point mutation in B-box-2 disrupted dimerization (Wagner et al., 2016). Inspired by this prior work, we designed, expressed, and purified two analogous miniTRIM37s comprising RING–B-box-2–hairpin coiled-coil (RBhcc) and B-box-2–hairpin coiled-coil (Bhcc) (**Fig. 4D**; **Fig, S4C**), and analyzed them by size exclusion chromatography coupled to multi-angle light scattering (SEC-MALS). Like the miniTRIM5α RBhcc, miniTRIM37 RBhcc was dimeric (**Fig. 4E**). However, in contrast to miniTRIM5α Bhcc (Wagner et al., 2016), miniTRIM37 Bhcc was monomeric (**Fig. 4E**). Thus, the B-box-2 of TRIM37 does not form an interface sufficiently robust to support oligomerization. Instead, these results suggest that oligomerization of TRIM37 dimers requires RING–RING interactions.

### The TRIM37 B-box 2 interface is dispensable for dimerization but impacts oligomer conformation

Consistent with our biochemical results, AlphaFold modeling of a TRIM37 RBhcc dimer revealed an extensive dimer interface between the two RING domains (**Fig. 5A**; **Fig. S4D**), with a predicted buried surface area per protomer of about 1070 Å^2^. The interface is predominantly hydrophobic, with I9, F13, L27, V64, V72, and L79 making symmetric contacts across the dimer interface (**Fig. 5B**); the predicted RING–RING interface almost perfectly matches a prior crystal structure of the TRIM37 RING domain dimer (PDB 3LRQ; <0.5 Å Cα rmsd in a single chain, and a near-exact dimer packing match). Despite being insufficient to mediate oligomerization, the B-box-2 domain in miniTRIM37 RBhcc is predicted to form a second, smaller, dimer interface that is structurally analogous to that in the TRIM5α B-box-2 dimer (**Fig. 5B**; (Wagner et al., 2016)). Three residues anchor the predicted B-box-2 interface: H115 makes symmetric pi-stacking interactions across the dimer interface, and L119 and W120 form additional hydrophobic interactions. The crystal structure of the TRIM5α B-box-2 dimer revealed a three-layer interface: Layer 1 involves electrostatic contacts between residues E102 and K103 across the dimer interface; Layer 2 involves hydrophobic and pi-stacking interactions between residues W117 and L118, and Layer 3 involves electrostatic interactions between residues E120, R121, and T130 (Wagner et al., 2016). The TRIM37 B-box interface has residues positioned similarly to one of the Layer 2 residues (H115 in TRIM37) and two of the Layer 3 residues (L119 and W120 in TRIM37).

**Figure 5.**
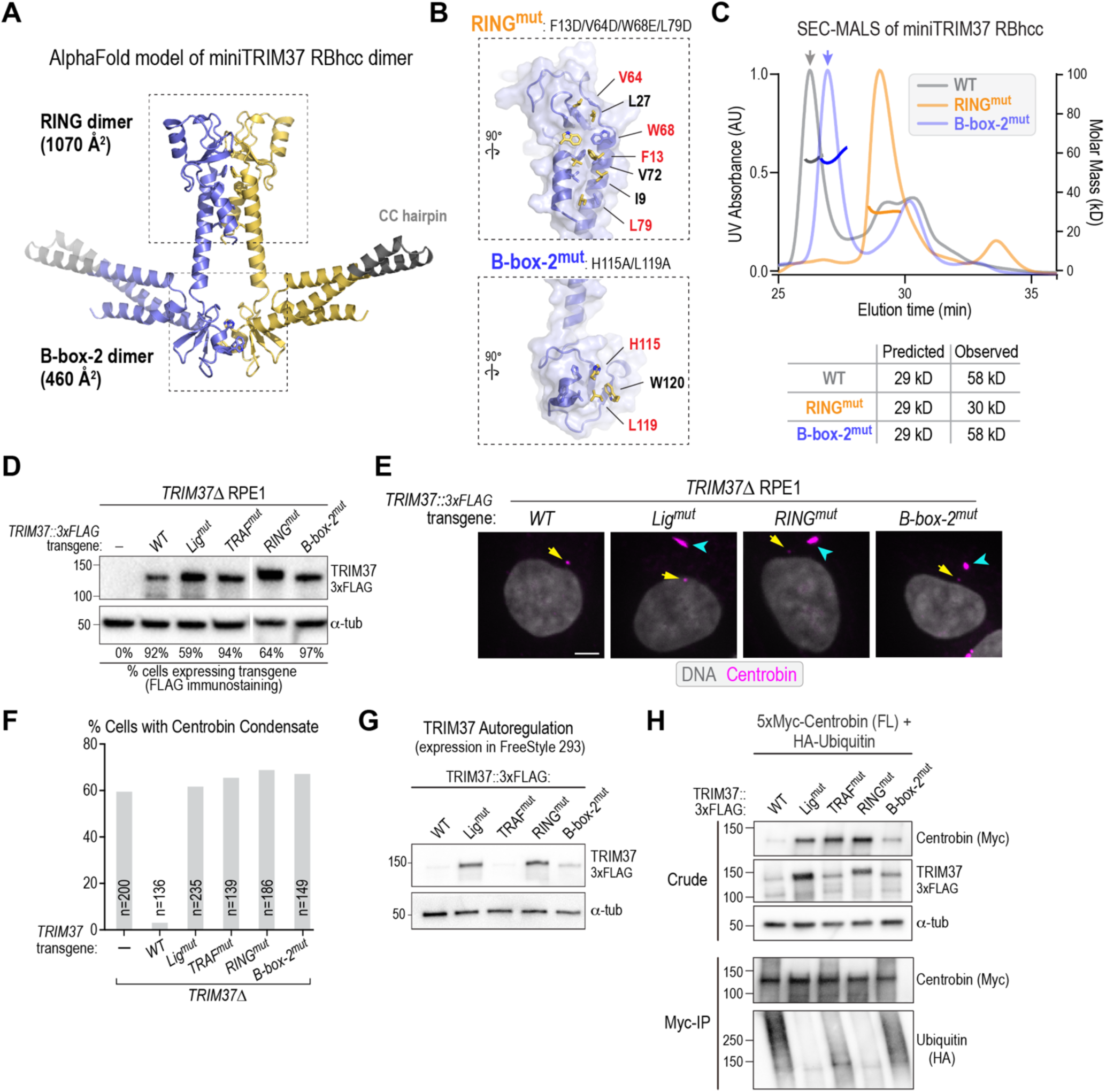
Functional analysis of predicted RING and B-box 2 interfaces of TRIM37. **(A)** AlphaFold model of the miniTRIM37 RBhcc dimer, highlighting the predicted RING-RING and B-box-2 – B-box-2 interfaces. **(B)** Detailed view of the 2 predicted interfaces, highlighting key residues. Mutations were engineered in the residues colored in red to disrupt key interface contact. **(C)** SEC-MALS analysis of the indicated miniTRIM37 RBhcc variants. Numbers next to the predicted molecular weights are the native molecular weights of the entities in the major UV peaks measured by MALS. The SEC-MALS analysis was repeated twice with identical results (see also *Fig. S4C*). The WT trace is reproduced from Fig. 4E for comparison. **(D)** Immunoblot of transgene-mediated expression of the indicated engineered TRIM37 variants in *TRIM37Δ* cells. The lanes shown are from the same gel and immunoblot; one intervening lane was removed, as indicated by the white line. α-tubulin serves as a loading control. The left four lanes are reproduced from Fig. 2B for comparison. **(E)** Immunofluorescence images of TRIM37Δ cells expressing the indicated transgene variants after fixing and staining for centrobin and DNA. Centrosomes (yellow arrows) and centrobin condensates (cyan arrowheads) are marked on the images. Scale bar is 5 µm. **(F)** Quantification of frequency of centrobin condensate formation for the indicated conditions. *n* is the number of cells analyzed. **(G)** Analysis of TRIM37 autoregulation following expression in FreeStyle 293F cells. Similar results were observed in two independent experiments. **(H)** Analysis of centrobin ubiquitination by the indicated TRIM37 variants. HA-ubiquitin was co-transfected along with FL centrobin and indicated TRIM37 variants. The ubiquitination assessed is on the small pool of soluble FL centrobin following immunoprecipitation. Similar results were observed in two independent experiments.

We designed two mutants to disrupt the predicted RING–RING and B-box-2–B-box-2 interfaces and assess their impact on miniTRIM37 RBhcc dimerization. For the RING interface, we mutated four key hydrophobic residues to charged residues: F13D, V64D, W68E, and L79D (**Fig. 5B**); these residues are far from the E2-binding interface (Gundogdu and Walden, 2019; Plechanovova et al., 2012) and are not expected to affect E2 binding. For the B-box-2 interface, we mutated both H115 and L119 to alanine (**Fig. 5B**). We then compared the two mutants to wildtype miniTRIM37 RBhcc using SEC-MALS (**Fig. 5C; Fig. S4C**). The results showed that mutation of the RING–RING interface disrupted dimer formation (**Fig. 5C**). By contrast, the B-box-2 interface mutant behaved as a homodimer, with a native molecular weight identical to wildtype miniTRIM37 RBhcc (**Fig. 5C**). These results confirm that TRIM37 cross-dimer interactions are primarily mediated by the RING domain. Interestingly, the miniTRIM37 RBhcc B-box-2 mutant reproducibly exhibited a shift to a later elution volume in size exclusion chromatography (**Fig. 5C**), which is consistent with the mutant dimer adopting a more compact conformation compared to the WT dimer. By contrast, when the same B-box-2 interface mutations were introduced into monomeric miniTRIM37 Bhcc, there was no difference in elution volume (**Fig. S4E**). Thus, while the B-box-2 interface is not required for oligomerization, it potentially affects the conformation of TRIM37 oligomers. These data highlight a significant difference between TRIM5α and TRIM37: the B-box-2 interface is critical for oligomerization of the former but dispensable for the latter, where it potentially impacts oligomer architecture.

### RING and B-box-2 interface mutants both compromise TRIM37 function

To test the functional importance of the RING and B-box-2 interfaces *in vivo*, we introduced transgenes expressing TRIM37 variants with mutations in *TRIM37Δ* RPE1 cells. After confirming expression (**Fig. 5D**), we compared the frequency of centrobin condensate formation. The results showed that both the RING and B-box-2 interfaces are important for TRIM37 to prevent centrobin condensate formation *in vivo* (**Fig. 5E,F**). Thus, while the B-box-2 interface is not required for oligomerization, it is critical for TRIM37 function.

We next monitored the impact of mutating the RING and B-box-2 interfaces on the autoregulation of TRIM37 by its ligase activity. Our prior work showed that the C18R mutation, which impairs ligase activity, stabilizes TRIM37 and significantly elevates its expression relative to WT TRIM37 (**Fig. 5G**; (Meitinger et al., 2020)). Similar to the ligase mutant, expression of the RING interface mutant was also elevated compared to WT TRIM37 (**Fig. 5G**). By contrast, the B-box-2 interface mutant had a minor effect, and the TRAF domain mutant had no effect (**Fig. 5G**). Thus, TRIM37 autoregulation relies on the RING interface and ligase activity but is largely independent of the B-box-2 interface and the TRAF domain.

Finally, we analyzed full-length centrobin ubiquitination by TRIM37 variants using co-expression analysis in human cells (**Fig. 5H**). Consistent with prior work, the results revealed robust ubiquitination by wild-type TRIM37 and no ubiquitination by the ligase mutant. The RING interface mutant behaved similarly to the ligase mutant. Ubiquitination was significantly reduced compared to wild-type TRIM37 by the B-box-2 interface mutant and even further reduced for the TRAF domain mutant (**Fig. 5H**). In support of this data, immunoblotting of crude cell extracts to assess centrobin levels, which reflects how efficiently it is targeted for ubiquitination by TRIM37, revealed significantly higher centrobin levels when co-expressed with the ligase-mutant, TRAF-mutant, and RING-mutant of TRIM37, relative to WT TRIM37. Centrobin levels were mildly elevated with the B-box-2 mutant of TRIM37 but not nearly to the same extent as the other mutants (**Fig. 5H**).

These results indicate that the combination of TRAF domain-mediated recognition of specific motifs, the RING oligomerization interface, and the B-box-2 interface collectively prevent the formation of centrobin condensates. Notably, the B-box-2 interface is dispensable for oligomerization and only modestly affects ubiquitin ligase activity. However, the B-box-2 interface appears to control the conformation of TRIM37 oligomers, which may be important for their ability to geometrically sense ectopic centrobin assemblies and trigger their ubiquitination-mediated clearance *in vivo*.

### Detection of centrobin condensates *in vivo* requires the TRIM37 B-box-2 interface

To address the role of the B-box-2 interface in localizing to centrobin condensates *in vivo*, we took advantage of the fact that ligase-mutant TRIM37 binds stably to the condensates but cannot disassemble them (Meitinger et al., 2021). We expressed mutated TRIM37 variants that combined the ligase mutation (C18R) with the TRAF mutant or with mutations that disrupted the B-box-2 interface or RING interface in *TRIM37Δ* cells and monitored their localization (**Fig. 6A**). We quantified the ratio of the TRIM37 variant, detected using an epitope tag on the transgene-encoded protein, to the centrobin signal at condensates. This analysis revealed that, while the ligase-mutant concentrated robustly on centrobin condensates, the ligase-and-TRAF double mutant failed to do so (**Fig. 6A-C**). Combining the ligase mutant with either the B-box-2 or RING interface mutants also compromised condensate localization (**Fig. 6A-C**). Consistent with this finding, the B-box-2 interface, RING interface, and TRAF single mutants were also poorly associated with centrobin condensates (**Fig. S5A,B**). These data indicate that the B-box-2 interface is critical for the recognition of oligomerized substrates *in vivo*. We suggest that the role of the B-box-2 interface is to geometrically sense centrobin oligomeric assemblies and target them for ubiquitination by stabilizing the substrate-bound active conformation of TRIM37 oligomers (**Fig. 6D**).

**Figure 6.**
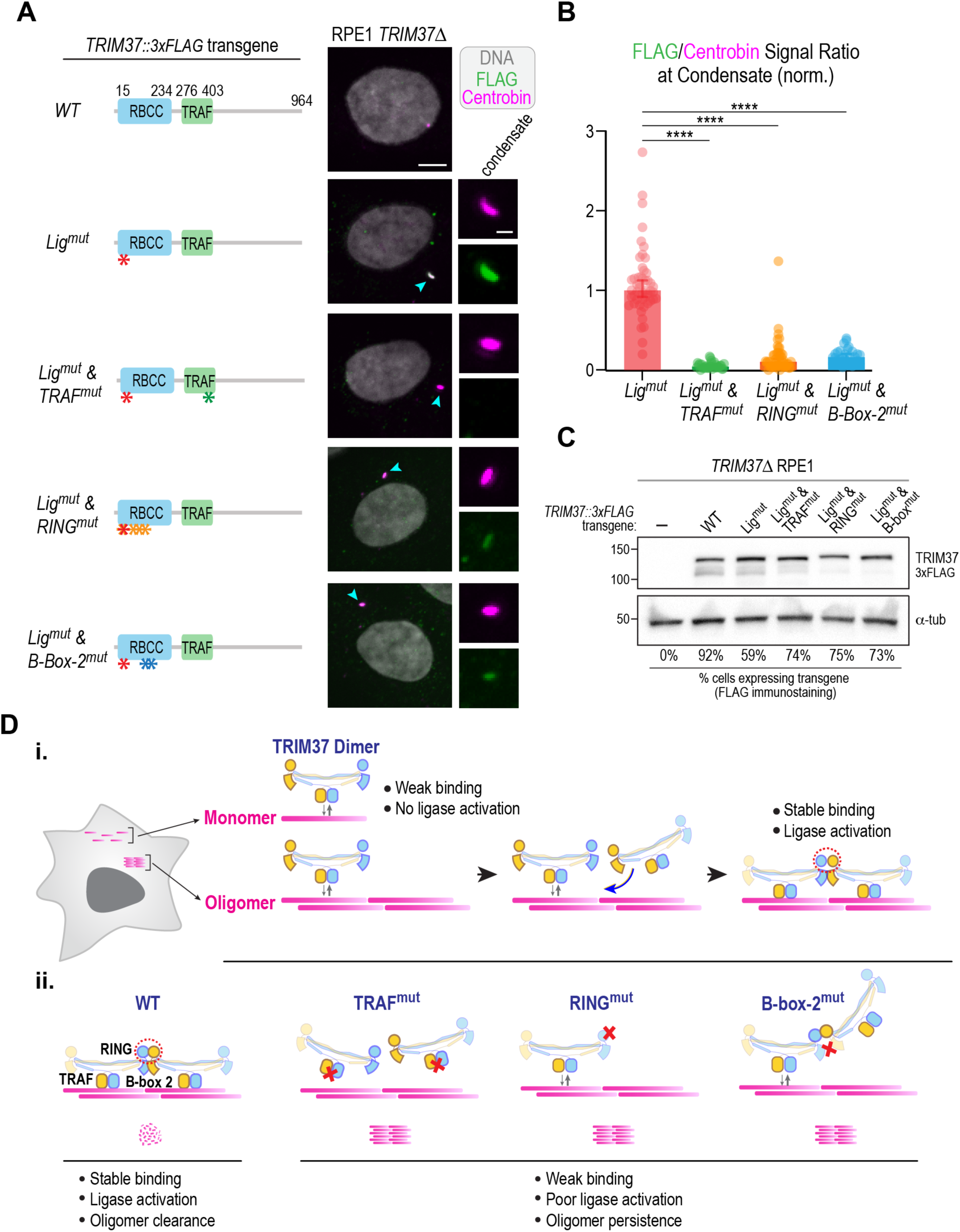
The RING and B-box-2 interfaces, along with the TRAF domain, facilitate detection of centrobin oligomers *in vivo*. **(A)** (*left*) Schematics highlighting TRIM37 variants expressed from transgenes in *TRIM37Δ* cells; (*right*) images of cells expressing the TRIM37 variants that were fixed and labeled for centrobin and for the FLAG epitope fused to the transgene-encoded TRIM37 variants. Condensates, when present, are highlighted by cyan arrowheads and are shown in a magnified view on the right, with separated centrobin (*magenta*) and FLAG (*green*) channels. **(B)** Quantification of the ratio of FLAG to centrobin signal at condensates for the indicated TRIM37 variants. The plotted values were normalized relative to the median value of the TRIM37 ligase-mutant. p-values are from unpaired t-tests; ****: p<0.001. **(C)** Immunoblot of transgene-mediated expression of the indicated engineered TRIM37 variants in *TRIM37Δ* cells. α-tubulin serves as a loading control. **(D)** Model for the substrate-dependent oligomerization and activation of TRIM37. Model highlights how TRIM37 selectively targets oligomers (i) and summarizes the effect of TRIM37 mutations (ii).

## DISCUSSION

Centrosomes are multi-gigadalton macromolecular assemblies consisting of ∼150 proteins organized into multiple substructures (Laporte et al., 2024; LeGuennec et al., 2021; Ma et al., 2023). Many of these interconnected structures (centriolar microtubules, cartwheel, inner scaffold, pericentriolar material) are polymeric. How the dimensions of these polymeric structures are specified and how ectopic polymeric assemblies containing centrosomal proteins are prevented from forming in the cytoplasm are important questions. While TRIM37 was initially thought to act at peroxisomes (Kallijarvi et al., 2002; Wang et al., 2017), recent work has suggested that its primary role is to restrict the growth of several ectopic polymeric assemblies of centrosomal proteins (Balestra et al., 2021; Meitinger et al., 2021; Meitinger et al., 2020). In this work, we address the molecular mechanism by which TRIM37 recognizes and prevents assembly of ectopic spindle poles that form on structured condensates of the centrosomal protein centrobin. These ectopic poles elevate chromosome missegregation during cell division and likely contribute to the high tumor incidence and other phenotypes in Mulibrey nanism patients (Balestra et al., 2021; Meitinger et al., 2021). Our prior work showed that deleting the gene encoding centrobin suppresses ectopic spindle poles in cells lacking TRIM37 function (Meitinger et al., 2021). Thus, a key question was how TRIM37 recognizes and selectively promotes the clearance of ectopic oligomeric centrobin condensates while not targeting centrobin monomers in the cytoplasm or endogenous centrobin assemblies at centrioles that are important for ciliogenesis (Karasu et al., 2022; Ogungbenro et al., 2018). Our results suggest a mechanism in which substrate-guided oligomerization of TRIM37 dimers increases the avidity of binding to ectopic assemblies and activates ligase activity to promote their ubiquitination and clearance (**Fig. 6D**).

### A mechanism for TRIM37-mediated selective suppression of centrobin oligomers

Formation of centrobin condensates in cells lacking TRIM37 occurs via the accumulation of material at centrosomes that eventually buds off to form an acentrosomal condensate. These condensates mature at an appreciable frequency into ectopic spindle poles that elevate chromosome missegregation and likely contribute to the tumor-prone nature of the *TRIM37* loss-of-function disorder Mulibrey nanism. Using the PLK4 inhibitor centrinone, together with an inducible *TRIM37* knockout, we show that centrosomes are not required for condensate formation in *TRIM37Δ* cells; condensates can initiate and expand, likely via centrobin oligomerization, without the need for centrosomes as a nucleating structure. Thus, TRIM37 can patrol the cytosol to detect and prevent the formation of ectopic centrobin-containing assemblies.

Using *in vivo* replacement of TRIM37 and centrobin, in combination with *in vitro* biochemical analysis and structural modeling, we propose a mechanism that enables TRIM37 to detect and clear nascent centrobin oligomers without targeting centrobin monomers that are recruited from the cytoplasm to function at centrioles and enable ciliogenesis (**Fig. 6D**). The first element required for the role of TRIM37 in this clearance is its TRAF domain, which follows the coiled-coil and is unique in the extended TRIM family. Consistent with the function of other TRAF domains, which exhibit binding specificity for different peptide motifs (Park, 2021; Zapata et al., 2007), our results implicate the TRIM37 TRAF domain in specific binding to peptide motifs in centrobin. Our results further show that disrupting either the peptide binding interface of the TRAF domain or the peptide motifs in centrobin prevents TRIM37 from detecting and clearing oligomeric centrobin condensates. Thus, analogous to the similarly positioned SPRY domain of TRIM5α, which binds the subunits of viral nucleocapsids in the cytosol (Li et al., 2016; Stremlau et al., 2006), the TRIM37 TRAF domain detects motifs in centrobin. Comparison of the *in vivo* localization of ligase-mutant TRIM37, which strongly concentrates on centrobin condensates, to that of ligase-and-TRAF-mutant TRIM37, which does not localize to centrobin condensates, provided strong support for the conclusion that the TRAF domain provides key specificity in the recognition of centrobin assemblies.

As TRAF domain-mediated motif recognition is low-affinity and would be unable to distinguish centrobin monomers from oligomers, additional elements of TRIM37 must also contribute to the specific recognition and clearance of centrobin oligomers. Based on the fact that TRIM family proteins are dimerized by an anti-parallel coiled-coil that places their two E3 ligase RING domains on opposite ends and that many RING ligases of the TRIM family must dimerize to fully activate the ligase (Fiorentini et al., 2020), we hypothesized that the oligomeric nature of the substrate facilitates the inter-dimer association of RING domains to promote ligase activation. In the case of TRIM5α, which employs such a mechanism (Ganser-Pornillos and Pornillos, 2019; Spada et al., 2024), the B-box-2 interface is critical for the inter-dimer association of RING domains and ligase activation (Wagner et al., 2016). By contrast, TRIM37 primarily employs a RING–RING interface for inter-dimer oligomerization, representing the second critical element required for TRIM37 dimers to clear ectopic centrobin assemblies. Notably, while the B-box-2 interface of TRIM37 is dispensable for inter-dimer oligomerization *in vitro*, its disruption resulted in a more compact conformation of the cross-dimer and prevented detection and clearance of centrobin condensates *in vivo*. Thus, the B-box-2 interface represents the third critical element required for TRIM37 to clear centrobin oligomers. We speculate that this requirement arises because TRIM37 dimers are present at relatively low concentrations *in vivo*, where the geometric properties that the B-box-2 interaction confers to the inter-dimer interface are required for efficient substrate-templated TRIM37 oligomerization (**Fig. 6D**). Thus, analogous to the way that trimerization of the TRIM5α B-box-2 is thought to match the lattice arrangement of nucleocapsid subunits to mediate viral recognition (Li et al., 2016; Wagner et al., 2016), the TRIM37 B-box-2 may function to recognize the specific geometry of ectopic centrobin assemblies. Our data suggest that TRAF domain-mediated substrate recognition, RING dimerization, and B-box-2-mediated sensing of oligomer geometry ensure that centrobin oligomers are selectively targeted for ubiquitination and degradation. Consistent with the importance of these TRIM37 domains in clearing centrobin condensates, we note that in addition to mutations that introduce early stop codons or delete the *TRIM37* gene, Mulibrey nanism patients have been identified with mutations in the B-box-2 (C108S) and with a 17 amino acid deletion in the TRAF domain (OMIM; https://www.omim.org/; (Amberger et al., 2015)).

In contrast to the requirements to clear centrobin oligomers, the most significant elements for TRIM37 to autoregulate its own levels are the RING interface and ligase activity; the TRAF interface is dispensable, and the B-box-2 interface only makes a mild contribution. Thus, at sufficient concentrations, the RING interface alone can activate the ligase and trigger self-destruction. By destroying TRIM37 that oligomerizes in the absence of substrate templating, this mechanism may serve to tune the concentration of TRIM37 to ensure that only ectopic centrobin oligomers, and not cytoplasmic centrobin or centriole-associated centrobin, become TRIM37 targets.

In addition to centrobin, the centrosomal protein CEP192 is a TRIM37 target and has received significant interest because its reduction following elevated TRIM37 expression due to genomic amplification in specific cancers leads to synthetic lethality with PLK4 inhibition (Meitinger et al., 2020; Yeow et al., 2020). TRIM37 also binds PLK4, can ubiquitinate it, and prevents the formation of PLK4-dependent foci that accelerate spindle formation following centrosome removal (Meitinger et al., 2016; Meitinger et al., 2021; Meitinger et al., 2020). Addressing how TRIM37 regulates these other targets will be influenced by the in-depth analysis of centrobin that we present here. We note that TRIM37 loss also leads to the formation of centrin-containing foci whose composition and functional significance are unclear (Balestra et al., 2021; Meitinger et al., 2021). While the key scaffold of these centrin-containing foci has not yet been defined, their existence suggests that TRIM37 has additional centrosomal protein substrates.

Both prior work and the analysis presented here cement the idea that TRIM37 acts as a guardian of the centrosome. Given its mutation in the human tumor-prone developmental disorder Mulibrey nanism and its elevation leading to synthetic lethality with PLK4 inhibition in specific cancers, the work presented here will aid in understanding the full spectrum of TRIM37 roles *in vivo*. An important open question is whether the types of assemblies formed when TRIM37 is absent are physiologically relevant in specific contexts where TRIM37 is downregulated, for example, as part of a developmental program. Here, we note that the *CNTROB* gene was identified due to a natural mutation in rats that impacts the formation of complex structures important for spermatogenesis (Liska et al., 2009), and centrobin was later shown to be important for ciliogenesis (Gottardo et al., 2015; Karasu et al., 2022; Ogungbenro et al., 2018; Reina et al., 2018). Selectively perturbing the oligomerization of TRIM37 substrates such as centrobin, independently of their control by TRIM37, may enable testing whether the assemblies observed in the absence of TRIM37 are physiologically significant.

## ACKNOWLEDGEMENTS

The authors thank Andrew Holland and Peter Yeow for communicating unpublished results and for discussion and Rebecca Green for help with the model figure. This work was supported by grants from the NIH to K.O. (R01 GM074207), A.D. (R01 GM074215), and K.D.C. (R35 GM144121). A. Brambila and F. Mendez-Diaz were IRACDA fellows and were supported by NIGMS/NIH K12 GM068524. K.O. acknowledges partial salary support from the Ludwig Institute for Cancer Research.

## AUTHOR CONTRIBUTIONS

Conceptualization: A. Desai, K.D.C., K.O.; Methodology: A. Bellaart, A. Brambila, J.X., F.M.D., A. Deep, J.A., K.D.C., A. Desai, K.O.; Formal analysis: A. Bellaart, A. Brambila, J.X., F.M.D., A. Deep; Investigation: A. Bellaart, A. Brambila, J.X., F.M.D., A. Deep., J.A., F.M., M.O.; Resources: A. Desai, K.D.C., K.O.; Writing - original draft: A. Desai, K.O.; Writing - review & editing: A. Bellaart, A. Brambila, J.X., F.M.D., A. Deep, F.M., M.O., K.D.C, A. Desai, K.O.; Visualization: A. Bellaart, A. Brambila, J.X., A. Deep, K.D.C, A. Desai, K.O.; Supervision: A. Desai, K.D.C., K.O.; Project administration: A. Desai, K.D.C., K.O.; Funding acquisition: A. Desai, K.D.C., K.O.

## SUPPLEMENTAL FIGURES WITH TITLES AND LEGENDS

**Figure S1.**
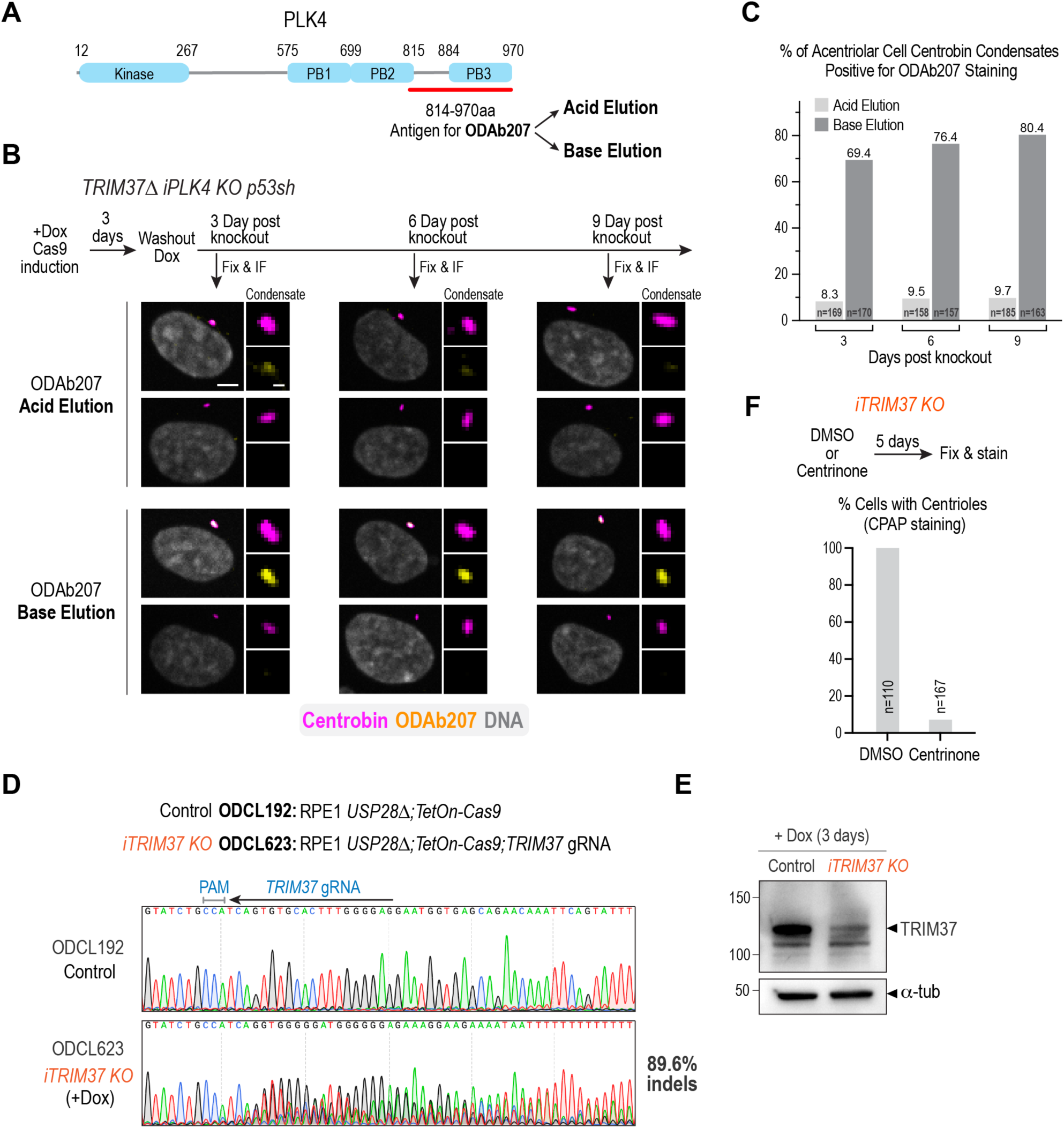
Analysis of PLK4 localization to centrobin condensates and validation of the inducible *TRIM37* knockout. **(A)** Schematic of PLK4 highlighting key domains and location of the antigen used to generate ODAb207 in rabbits. Following serum recirculation on the antigen column, antibodies were eluted first using low pH (Acid eluction) and second using high pH (Base elution). The pH of eluted fractions was rapidly neutralized and the Acid and Base elutions were separately dialyzed into a storage buffer. **(B)** Analysis of ODAb207 staining following inducible knockout of PLK4 in *TRIM37Δ p53sh* RPE1 cells. Knockout efficiency was confirmed by the absence of centrioles (*not shown*). Two examples are shown for the Acid and Base ODAb207 elutions. For each elution, the top example shows labeling (strong or weak) with ODAb207 of condensates marked by centrobin, while the bottom example shows no labeling. The Base but not the Acid elution of ODAb207 robustly labeled centrobin condensates. Scale bars, 5 µm and 1 µm. **(C)** Graphical summary of ODAb207 staining of centrobin condensates. Only condensates in acentriolar cells, indicative of *PLK4* knockout, were analyzed. Condensate labeling was prominent with the Base elution of ODAb207, even 9 days after knockout induction; by contrast, little-to-no labeling was observed with the Acid elution of ODAb207. *n* is the number of cells analyzed. The analysis of the inducible PLK4 knockout and the predominant labeling of centrobin-containing condensates with the Base but not Acid elution of ODAb207 indicate that the observed staining does not represent PLK4 and instead represents cross-reactivity with a condensate component. **(D)** Sequencing traces of control (ODCL192) and *iTRIM37 KO* (ODCL623) cells following 3 days of doxycycline induction of Cas9. The sequence of the *TRIM37* gRNA is indicated above the control sequencing trace. The frequency of indels was estimated using TIDE analysis (see Methods) and is derived by subtracting the percentage of 0 bp changes (10.4%) from 100%. **(E)** Immunoblot of the cell lines shown in *(D)* highlighting significant reduction in TRIM37 protein after knockout induction. α-tubulin serves as a loading control. **(F)** Quantification of centriole depletion following 5-day centrinone treatment of *iTRIM37 KO* cells. Centrinone was used to first deplete centrioles prior to inducing *TRIM37* knockout. *n* is the number of cells analyzed.

**Figure S2.**
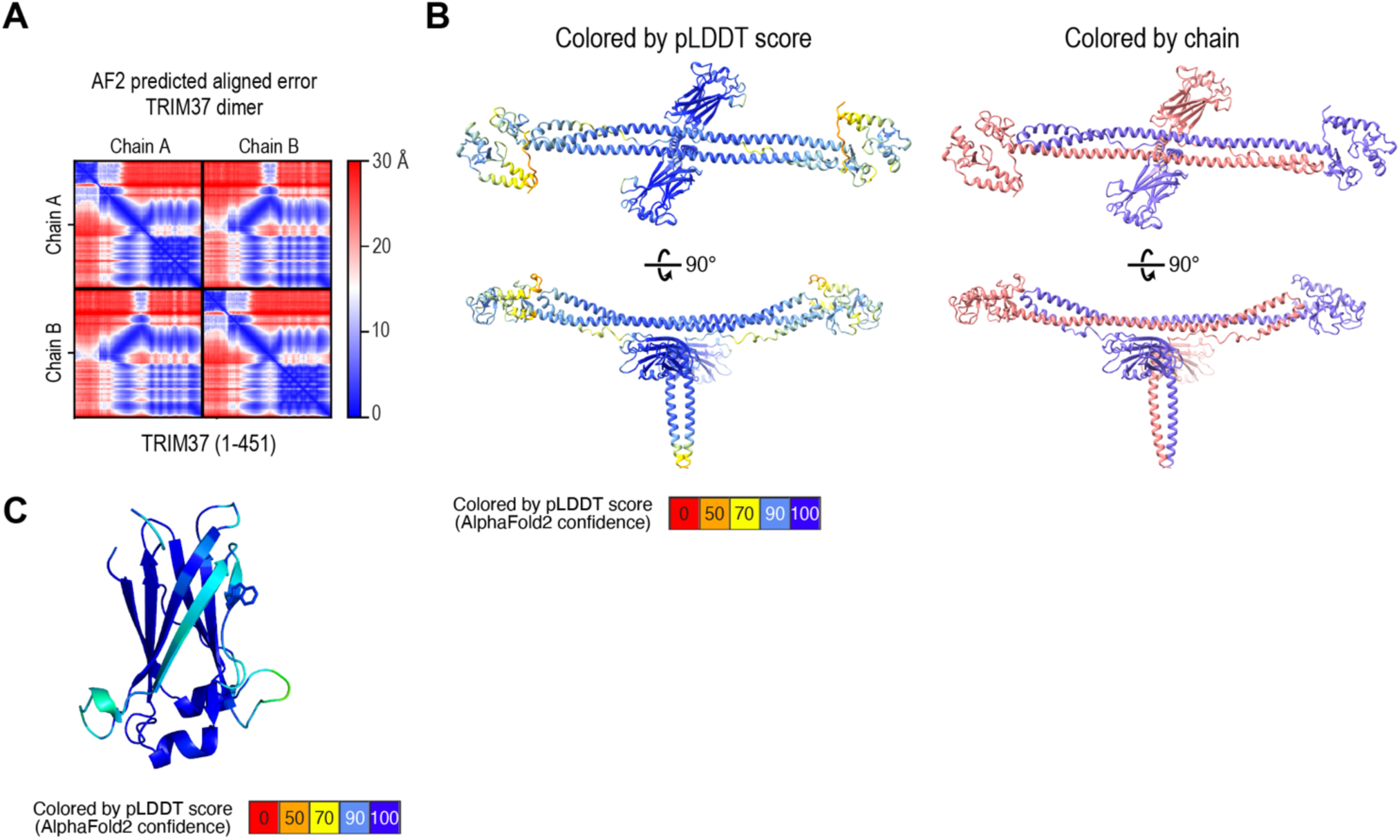
Error plots of Alphafold models of TRIM37 dimer and TRAF domain. **(A)** Predicted Aligned Error (PAE) plot of the TRIM37 dimer model. **(B)** TRIM37 dimer model colored by confidence (pLDDT score; *left*) and by chain (*right*). **(C)** TRIM37 TRAF domain model colored by confidence (pLDDT score).

**Figure S3.**
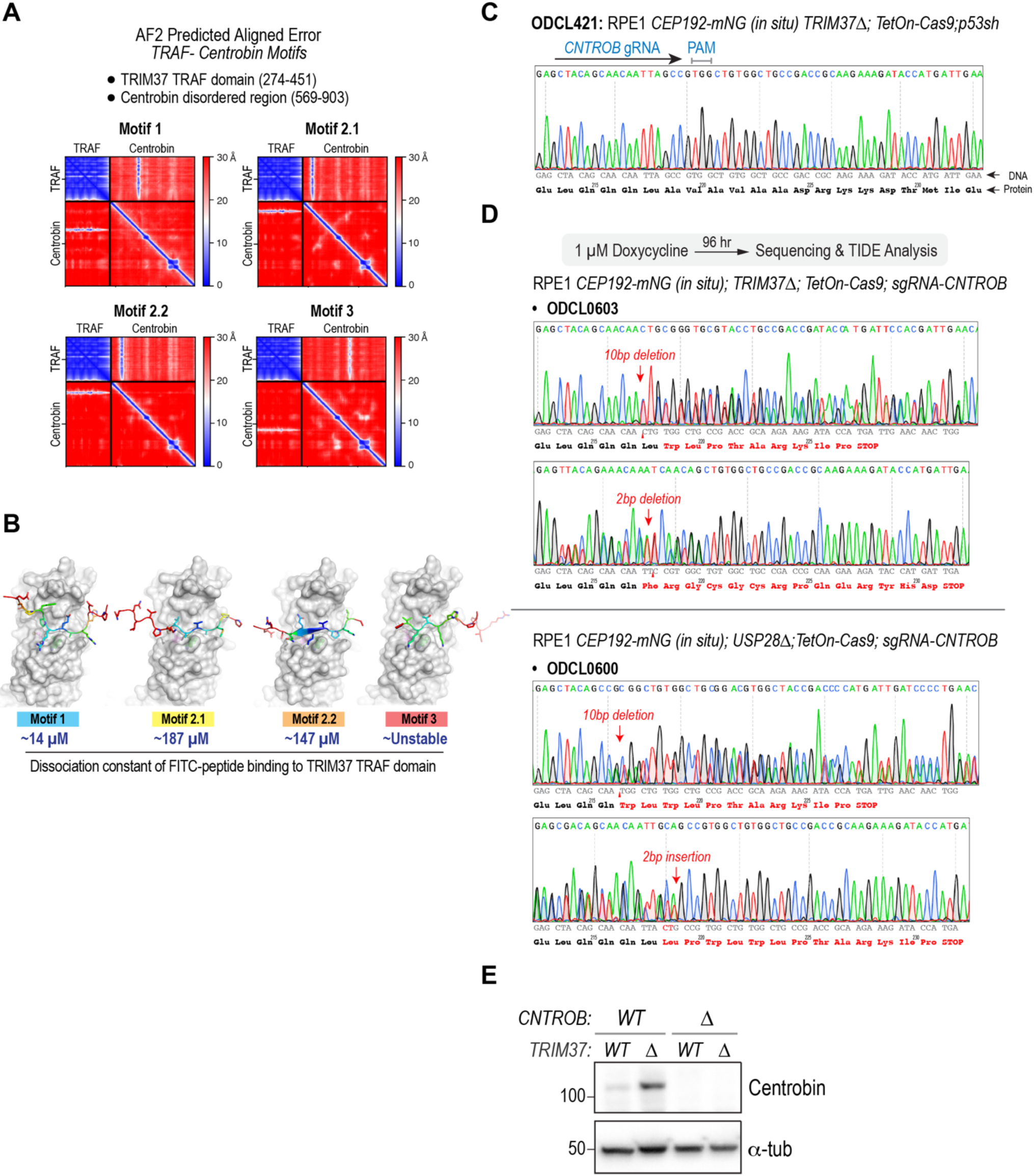
Error plots of TRAF-centrobin motif Alphafold models and validation of *CNTROB* knockouts. **(A)** PAE plots of TRIM37 TRAF–centrobin motif models. The regions of TRIM37 and centrobin used to generate the models are noted above. **(B)** TRAF-centrobin motif models (*same as in* Fig. 3B) with binding affinities of FITC-coupled motif peptides estimated from the curves shown in Fig. 3D. **(C)** Sequence trace of a control cell line highlighting the *CNTROB* gRNA. **(D)** Sequencing traces of *CNTROBΔ* cell lines with *TRIM37 WT* or *TRIM37*Δ. For each cell line, the top trace is with a forward sequencing primer and the bottom trace with a reverse sequencing primer (presented as the complementary strand). The changes in each allele are annotated on the traces. **(E)** Immunoblot verifying loss of centrobin protein in *CNTROBΔ* cells, with or without TRIM37 present. α-tubulin serves as a loading control.

**Figure S4.**
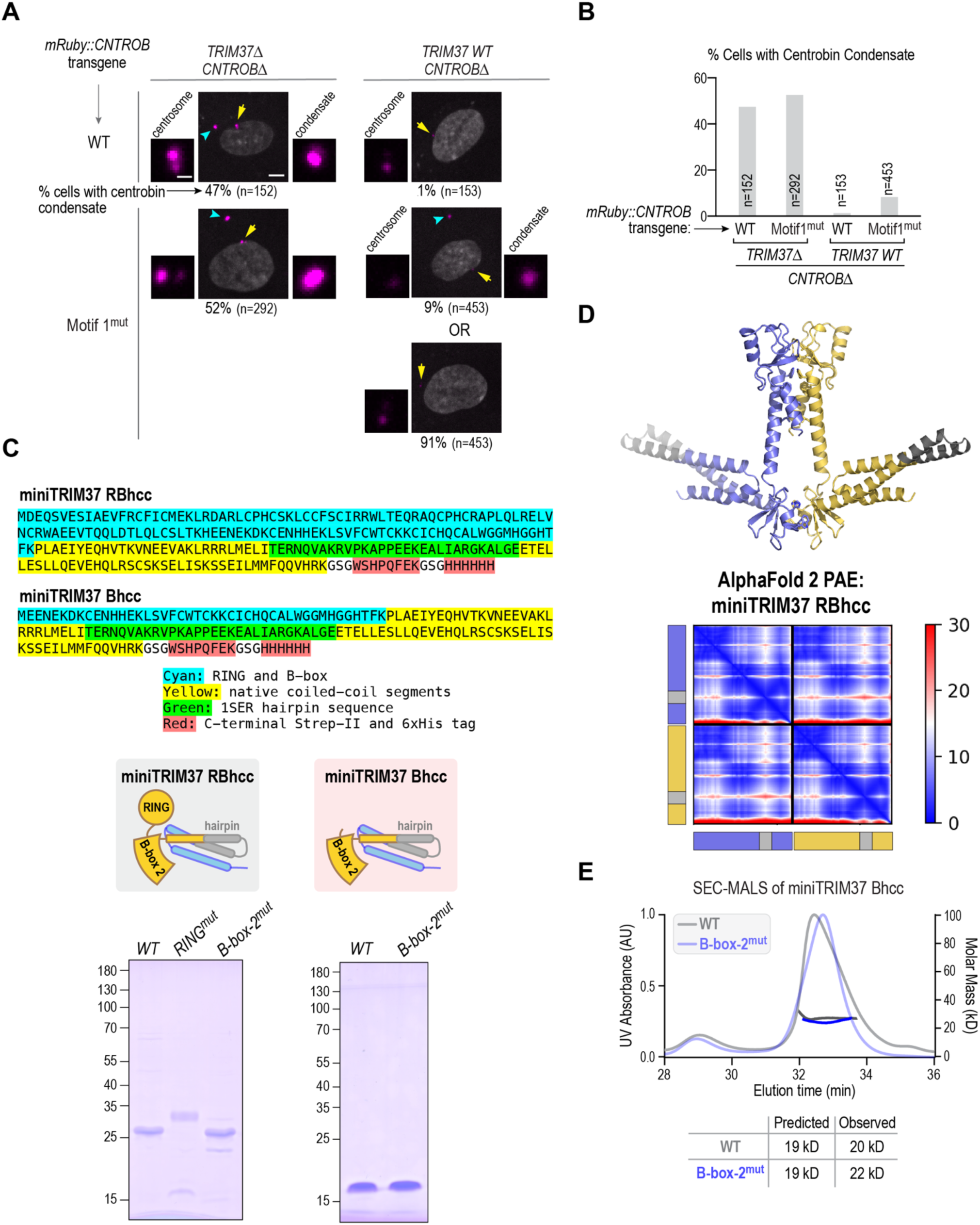
Analysis of Motif 1-mutant centrobin and of miniTRIM37s. **(A)** Images of *CNTROBΔ* cells, either without or with TRIM37 present, in which either mRuby-tagged WT or Motif 1-mutant centrobin was expressed from a lentivirally delivered transgene. For TRIM37 WT cells expressing the Motif 1-mutant, example images are shown of cells with and without a condensate. Scale bars, 5 µm and 1 µm. **(B)** Quantification of centrobin condensate frequency for the indicated conditions. **(C) (***top*) Protein sequences of miniTRIM37 RBhcc and Bhcc. (*bottom*) Coomassie-stained gels of purified recombinant miniTRIM37 RBhcc and Bhcc proteins (schematized above the gels; the same schematics are shown in Fig. 4D). The RING mutant migrates higher than the wild-type protein, likely due to the negative charges introduced by the mutations made to disrupt the RING-RING interface. The identity of the RING mutant band was confirmed using mass spectrometry. **(D)** (*top*) miniTRIM37 RBhcc dimer model, also shown in Fig. 5A, and (*bottom*) PAE plot of the miniTRIM37 RBhcc dimer model. The color-coded two chains used to construct the model are shown on the left and bottom sides of the PAE plot. **(E)** SEC-MALS analysis of miniTRIM37 Bhcc WT and mutant purified proteins. Both proteins elute at the same time from the column and have the same native molecular weight, corresponding to the predicted monomer molecular weight.

**Figure S5.**
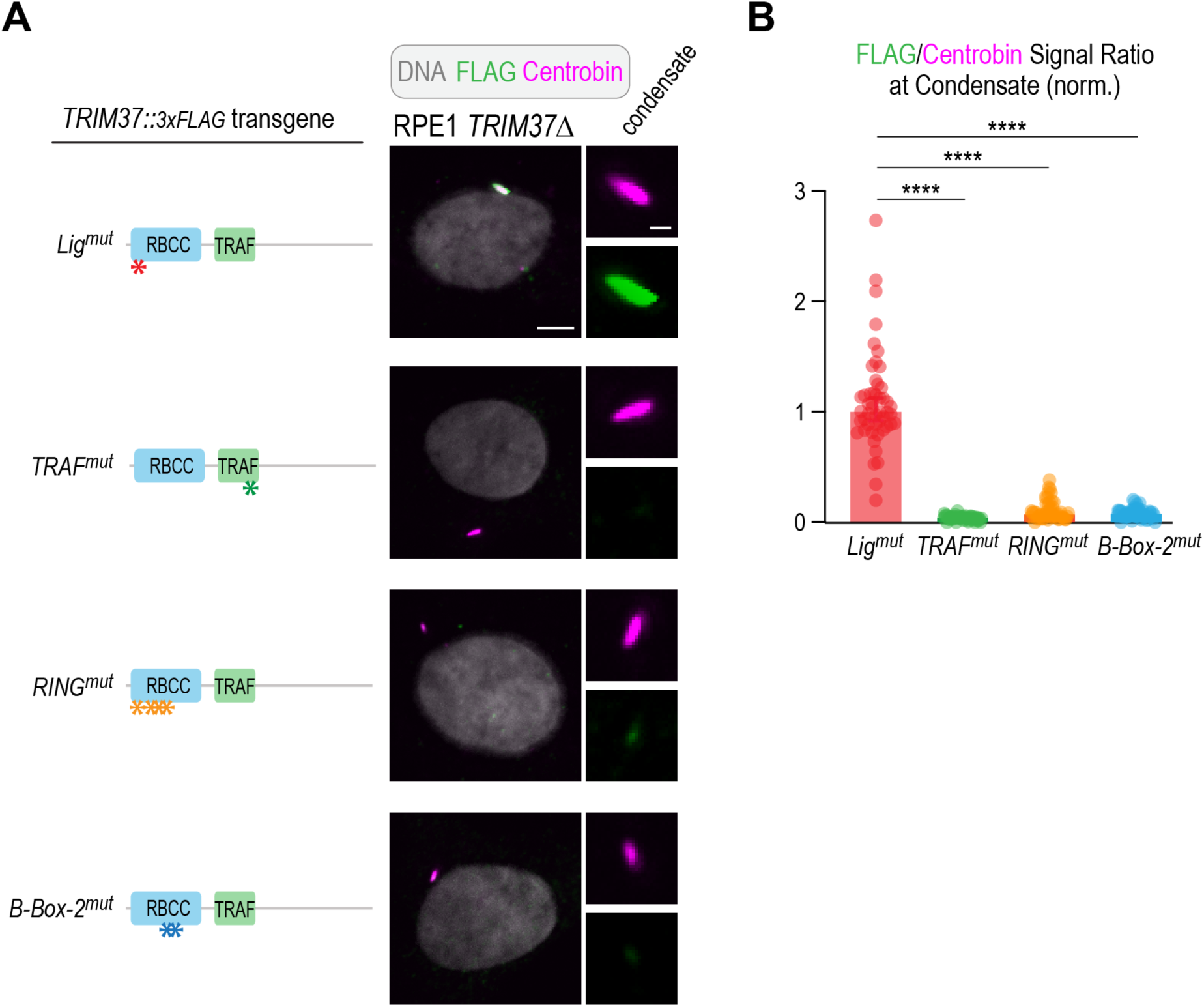
Analysis of TRIM37 single mutants localization to centrobin condensates. **(A)** (*left*) Schematics of transgene-encoded TRIM37::3xFLAG variants and (*right*) images of *TRIM37Δ* cells expressing indicated variants and labeled for centrobin and the FLAG epitope. Scale bars, 5 µm & 1 µm. **(B)** Quantification of the ratio of FLAG to centrobin signal at condensates for the indicated TRIM37 variants. The plotted values were normalized relative to the median value of the TRIM37 ligase-mutant. The data shown for the ligase-mutant is the same as in Fig. 6B. p-values are from unpaired t-tests; ****: p<0.0001.

## MATERIALS AND METHODS

### Cell line construction and culture

Cell lines used in this study are listed in **Table S1**. RPE1 cells were grown in F12/DMEM with 10% FBS, 100 µg/ml streptomycin, and 100 µg/ml penicillin. Lenti-X 293T cells were grown in DMEM with 10% tet system approved FBS (Thermo Fisher), 100 µg/ml streptomycin, and 100 µg/ml penicillin. Freestyle 293F cells were grown in Freestyle 293 expression medium (Thermo Fisher). Cells were cultured at 37°C in 5% CO_2_ except for Freestyle 293-F cells, which were cultured at 37°C in 8% CO_2_.

The RPE1 inducible knockout of *TRIM37* was generated by sequential lentiviral integration of inducible Cas9 (Edit-R Inducible Lentiviral Cas9; Dharmacon) and of a gRNA targeting exon 5 of *TRIM37* (pOD4913; 5’-TCAGTGTGCACTTTGGGGAG-3’). Cas9 expression was induced by adding doxycycline to a final concentration of 1 µg/ml and the efficiency of indel generation at the *TRIM37* locus was assessed using TIDE (Brinkman et al., 2014). To deplete centrosomes via PLK4 inhibition, centrinone was added at a final concentration of 100 or 150 nM.

RPE1 cell lines with *CNTROB* deleted were generated by treating the parent cell lines ODCL188 (*TRIM37 WT; USP28Δ*) and ODCL189 (*TRIM37Δ*), which contained inducible Cas9 and a gRNA targeting exon 5 of *CNTROB* (5’-GCTACAGCAACAATTAGCCG-3’), with doxycycline. Using limiting dilution the clonal cell lines ODCL0600 (*CNTROB*Δ; *TRIM37 WT; USP28Δ*) and ODCL0603 (*CNTROB*Δ; *TRIM37Δ*), which were confirmed by immunolabeling, immunoblotting, and genomic sequencing, were obtained. To generate the polyclonal inducible *PLK4 KO* cell line with p53 knocked down (ODCL420), a shRNA targeting p53 (pOD4894) was stably integrated using lentiviral delivery and the cell population was selected using the MDM2 inhibitor RG-7112 (Selleckchem).

Cell lines containing stably integrated FLAG-tagged *TRIM37* transgenes (wild-type, C18R, W373A, Ring^Mut^, B-Box-2^Mut^, C18R-W373A, C18R-Ring^Mut^, C18R-B-Box-2^Mut^, and Ring^Mut^-B-Box-2^Mut^; see **Table S1**) were generated by lentiviral delivery into the parent cell line ODCL61 (*TRIM37Δ*). Cell lines containing stably integrated transgenes encoding mRuby-centrobin (wild-type, TBM^Mut^, Motif-1^Mut^; see **Table S1**) were generated by lentiviral delivery into the parent cell lines ODCL600 (*CNTROB*Δ; *USP28*Δ) and ODCL602 (*CNTROB*Δ; *TRIM37*Δ).

For all lentiviral integrations, viral particles were prepared by transfecting the lentiviral construct into Lenti-X 293T cells using Lenti-X^TM^ Packaging Single Shots (Takara Bio USA, 631276). 72 hours after transfection, the virus-containing culture supernatant was collected and added to the growth medium of the cells to be transfected in combination with addition of polybrene (EMD Millipore) to 8 μg/ml. Polyclonal cell lines were selected using appropriate antibiotics (neomycin, 400 μg ml^−1^; puromycin, 10 μg ml^−1^ for RPE1 cells).

### Expression in Freestyle 293-F cells

For immunoprecipitation assays, plasmids containing centrobin constructs with a 5xMyc tag and TRIM37 constructs with a 3xFLAG tag were transfected into FreeStyle 293-F cells (Thermo Fisher) using FreeStyle MAX Reagent and OptiPRO SFM according to manufacturer guidelines (Thermo Fisher). Each transfected sample was a 20 ml culture (10^6^ cells/ml). Equal amounts (12.5 µg) of each expression plasmid were used for co-transfections. Following transfection, FreeStyle 293-F cells were incubated for 48 hours on an orbital shaker platform (125 rpm) at 37°C in 8% CO_2_. 10 ml of each sample were collected and washed with Dulbecco’s phosphate buffered saline (DPBS; Thermo Fisher). Cells were collected by pelleting and resuspended in 1 ml of lysis buffer (20 mM Tris/HCl, pH 7.5, 150 mM NaCl, 1% Triton X-100, 5 mM EGTA, 1 mM DTT, 2 mM MgCl and one EDTA-free protease inhibitor cocktail tablet (Roche). Cells were sonicated in a water bath sonicator at 4°C for 6 minutes to generate a crude lysate. The crude lysate was centrifuged at 13,000 rpm for 15 minutes at 4°C in a microfuge to generate supernatant and pellet fractions. Immunoprecipitations were performed by adding 20 µl of Pierce anti-Myc magnetic beads to 1 ml of supernatant (see **Table S3**). After incubating for 2 hours on a rotator at 4°C, beads were washed 5X in 900µl of lysis buffer and resuspended in 60 µl of 4x Laemmli sample buffer. For coimmunoprecipitation assays detecting ubiquitination, FreeStyle 293-F cells were transfected and incubated with equal amounts (8.5 µg) of three DNA constructs encoding: Myc-centrobin, FLAG-TRIM37, and haemagglutinin (HA)-tagged ubiquitin. For these IPs 5 mM N-ethylmaleimide was added to the lysis buffer.

### Immunofluorescence analysis

For immunofluorescence, 8,000 - 10,000 cells were seeded per well into 96-well plates one day prior to fixation. Cells were fixed in 100 μl of −20 °C methanol for 7 min. Cells were washed three times with wash buffer (phosphate-buffered saline (PBS) containing 0.1% Triton X-100) and blocked with blocking buffer (wash buffer containing 2% bovine serum albumin (BSA) and 0.1% sodium azide) overnight at 4°C or for 2 hours at RT. After blocking, cells were incubated for 2h with primary antibody (see **Table S3**) in blocking buffer at room temperature, followed by three washes with wash buffer. Cells were incubated for 1.5-2 hours at RT with the secondary antibody, stained with Hoechst 33342 dye in blocking buffer, and washed three times with wash buffer before imaging.

A directly labeled mouse polyclonal anti-centrobin antibody (see **Table S3**) was generated by using a Mix-n-Stain^TM^ CF568 dye antibody labeling kit (Biotium). Prior to adding the directly-labeled antibody, cells were blocked with normal mouse serum (Jackson Immunoresearch; 1:20 in wash buffer) for 2h, after which cells were incubated at room temperature with the directly-labeled antibody at a final concentration of 0.5 µg/ml in blocking buffer.

Images were acquired with a CQ1 spinning disk confocal system (Yokogawa Electric) equipped with a 40X (numerical aperture (NA) 0.95) U-PlanApo objective and a 2,560 × 2,160 pixel sCMOS camera (Andor). Image acquisition and data analysis were performed using CQ1 software and ImageJ, respectively. Images were also acquired on a Nikon ECLIPSE Ti2 spinning disk confocal system equipped with 40X dry (NA 0.95) or a 60X oil (NA 1.42) objectives and a 1,024 × 1,024 pixel iXon 888 EMCCD camera (Andor). Image acquisition and data analysis were performed using Image J and Nikon NIS-Elements software.

### Live-cell imaging

Samples were prepared 24 hours prior to imaging by seeding 4,000 cells per well into 96-well polystyrene plates. SiR-DNA was added for two hours prior to imaging at a final concentration of 0.5 µM per well. Live-cell imaging was performed with a CQ1 spinning-disk confocal microscope (Yokogawa Electric Corporation), using a 40X 0.95 NA U-Plan Apo objective at 37°C and 5% CO_2_. For imaging, 5 x 2 µm Z sections were acquired in 5 fields every 6 minutes for 14 hours in the following channels: far red/SiR-DNA (20% laser power, 200 ms exposure), red/mRuby::centrobin (50% laser power, 200 ms exposure), green/CEP192-mNG (20% laser power, 200 ms exposure).

### Quantification of centrobin and TRIM37::3xFLAG at condensates

Integrated fluorescence intensity was measured in a box fit around the condensate in the centrobin (Red) channel; the same box was transferred to the FLAG (far red) channel. The local per-pixel background was measured in a 1-pixel wide box around the selection. The normalized ratio of FLAG-to-centrobin signal was calculated by dividing the background-subtracted FLAG signal by the background-subtracted centrobin signal and dividing this ratio by the median value of the same ratio for ligase-mutant TRIM37.

### Design and construction of miniTRIM37 RBhcc and Bhcc expression plasmids

miniTRIM37 RBhcc and Bhcc constructs were designed using a similar strategy to the one employed previously for TRIM5α (Wagner et al., 2016). Specifically, miniTRIM37 RBhcc was designed by fusing residues 1-158 of human TRIM37 (containing the RING and B-box-2 domains, followed by a short segment of coiled-coil) to a serine tRNA synthetase hairpin sequence (resides 549-578, extracted from PDB 1D 1SER), followed by residues 214-254 of TRIM37; the deletion of residues 159-213 removes a majority of the antiparallel coiled coil. The annotated sequence of miniTRIM37 RBhcc is shown in *Fig. S4C*. WT, RING-mutant (F13D, V64D, W68E, and L79D) and B-box-2 mutant (H115A and L119D) miniTRIM37 RBhcc gBlock DNA sequences were obtained from IDT technologies. gBlocks contained extensions for ligation-independent V2 cloning (https://qb3.berkeley.edu/facility/qb3-macrolab/projects/lic-cloning-protocol/). For miniTRIM37 Bhcc constructs, PCR amplification was performed using the RBhcc gblocks as template DNA to add a start codon and delete the first 87 amino acids using the following oligos: Forward 5’ *<u>TTTAAGAAGGAGATATAGATCATG</u>*GAGGAGAACGAGAAAGATAAGTGC 3’ and reverse: 5’*<u>TTATGGAGTTGGGATCTTATTA</u>*GTGGTGATGGTGGTGGTGACC3’ (underlined sequences are for ligation-independent cloning). The annotated sequences of miniTRIM37 Bhcc is shown in *Fig. S4C*. miniTRIM37 RBhcc gblocks and Bhcc PCR-amplified DNA sequences were cloned into the pLICTr-NTA vector utilizing ligation-independent cloning and verified using sequencing. The AlphaFold dimer prediction of the engineered miniTRIM37 RBhcc is shown in *Fig. 5A*.

### Expression and Purification of miniTRIM37 RBhcc and Bhcc Proteins

miniTRIM37 RBhcc and Bhcc plasmids were transformed into Rosetta (DE3) *E. coli* cells employing a standard bacterial transformation protocol. For expression of each variant, 2 liters of 2XYT growth medium supplemented with 50 μM Zinc acetate were inoculated with overnight starter culture grown at 37°C in carbenicillin and chloramphenicol. Cultures were incubated at 37°C until the OD_600_ reached ∼0.9-1.0. Cultures were cooled on ice for 10 minutes to lower temperature and protein expression was induced by adding 1 mM IPTG. Induced cultures were incubated at 20°C for 4 hours, harvested and lysed via sonication in resuspension buffer (20 mM Tris-HCl pH 7.5, 300 mM NaCl, 10 mM Imidazole, 2 mM β-mercaptoethanol, and 10% glycerol) supplemented with a protease inhibitor mix (PMSF, leupeptin, pepstatin, and aprotinin). Lysates were clarified by centrifugation at 17,000 rpm at 4°C and supernatants loaded onto a 2 mL Ni-NTA column, washed with 10 column volumes of washing buffer (20 mM Tris-HCl pH 7.5, 300 mM NaCl, 20 mM Imidazole, and 10% glycerol), and eluted in 20 mM Tris-HCl pH 7.5, 150 mM NaCl, 400 mM Imidazole, and 10% glycerol. Eluates were concentrated at 4°C using Amicron Ultra spinning concentrators (10 kDa MW cutoff) and fractionated on a Superdex 200 gel filtration column in 50 mM Tris-HCl pH 8, 150 mM NaCl, 1 mM DTT, and 1 mM sodium azide. Protein purity was assessed via SDS-PAGE and fractions were concentrated to 1 mg/mL for SEC-MALS analysis.

### SEC-MALS analysis of miniTRIM37 RBhcc and Bhcc proteins

Prior to analysis of purified proteins, a Superdex 200 column coupled to a MALS machine (Wyatt Technology) was equilibrated overnight at room temperature in the buffer used for gel filtration. 120 µL of each purified protein (∼1 mg/ml) was auto-loaded onto the Superdex 200 column. Elution and light scattering of miniTRIM37 RBhcc and Bhcc proteins were monitored using Wyatt Technology HPLC software. After each run, UV absorbance and light scattering baselines were established, and protein peaks were defined to obtain the predicted molecular weights. SEC-MALS data was exported and graphed using GraphPad Prism software.

### Purification of recombinant TRIM37 TRAF domain

The TRIM37 TRAF domain coding sequence (residues 274-407) was amplified and cloned into the UC Berkeley Macrolab vector 2CT (Addgene number: 29706) to express N-terminal TEV protease-cleavable His_6_-MBP-tagged fusions. The TRAF W373A point mutant (TRAF^mut^) was generated using PCR-based site-directed mutagenesis, and constructs were verified by sequencing.

The MBP-TRIM37 TRAF and MBP-TRIM37 TRAF^mut^ expression constructs were transformed, and proteins were expressed in *E. coli* Rosetta2 pLysS (EMD Millipore). The cells were grown to an OD_600_ of 0.6-0.8, followed by induction with 0.33 mM IPTG. Protein expression was carried out at 20°C for 16-18 hours. Cells were harvested by centrifugation and resuspended in ice-cold resuspension buffer (50 mM Tris, pH 7.5, 300 mM NaCl, 10 mM imidazole, 10% glycerol, 2 mM β-mercaptoethanol). Resuspended cells were lysed using sonication, and the lysate was clarified by centrifugation. Proteins were purified by nickel affinity chromatography, and the eluted proteins were concentrated and further purified using size exclusion chromatography (Superdex 200 Increase 10/300 GL, Cytiva) in SEC buffer (20 mM Tris, 150 mM NaCl, and 1 mM DTT). Peak fractions were pooled and concentrated for use in binding assays.

### Fluorescence polarization-based peptide motif binding analysis

FITC-Ahx-labeled centrobin peptides, whose sequences are shown in *Fig. 3D*, were synthesized (BioMatik) and resuspended in DMSO at a concentration of 5 mM. Peptides were diluted in binding buffer (20 mM Tris, pH 7.4, 150 mM NaCl, 1 mM DTT) to a final concentration of 100 nM. Each 20 μL reaction mixture contained 100 nM peptide and specified varying concentrations of TRIM37 TRAF or TRIM37 TRAF^mut^ in the binding buffer. The reaction mixtures were incubated at room temperature for 20 minutes, and fluorescence polarization was measured using a TECAN Infinite M1000 PRO fluorescence plate reader in 384-well plates. The measurements were performed in triplicate, and the binding data were analyzed with GraphPad Prism v9 using a single-site binding model.

### AlphaFold2 modeling

Protein structure and interaction predictions were performed using the AlphaFold2 ColabFold notebook (Jumper et al., 2021; Mirdita et al., 2022). Residues 1-451 from TRIM37 were used to predict the homodimeric model shown in *Fig. 2A*. For predicting the interaction between TRIM37 TRAF and Centrobin, the TRIM37 TRAF domain (residues: 274-451) and Centrobin (residues: 569-903) were used in AlphaFold2. Structural analysis and visualizations were performed using PyMol (DeLano, 2002) and ChimeraX (Meng et al., 2023). The predicted buried surface area in the AlphaFold model of the TRIM37 RBhcc dimer was calculated using PDBePISA (https://www.ebi.ac.uk/pdbe/pisa/papers/pisa-web.pdf).

## SUPPLEMENTAL TABLES

**Table S1.**
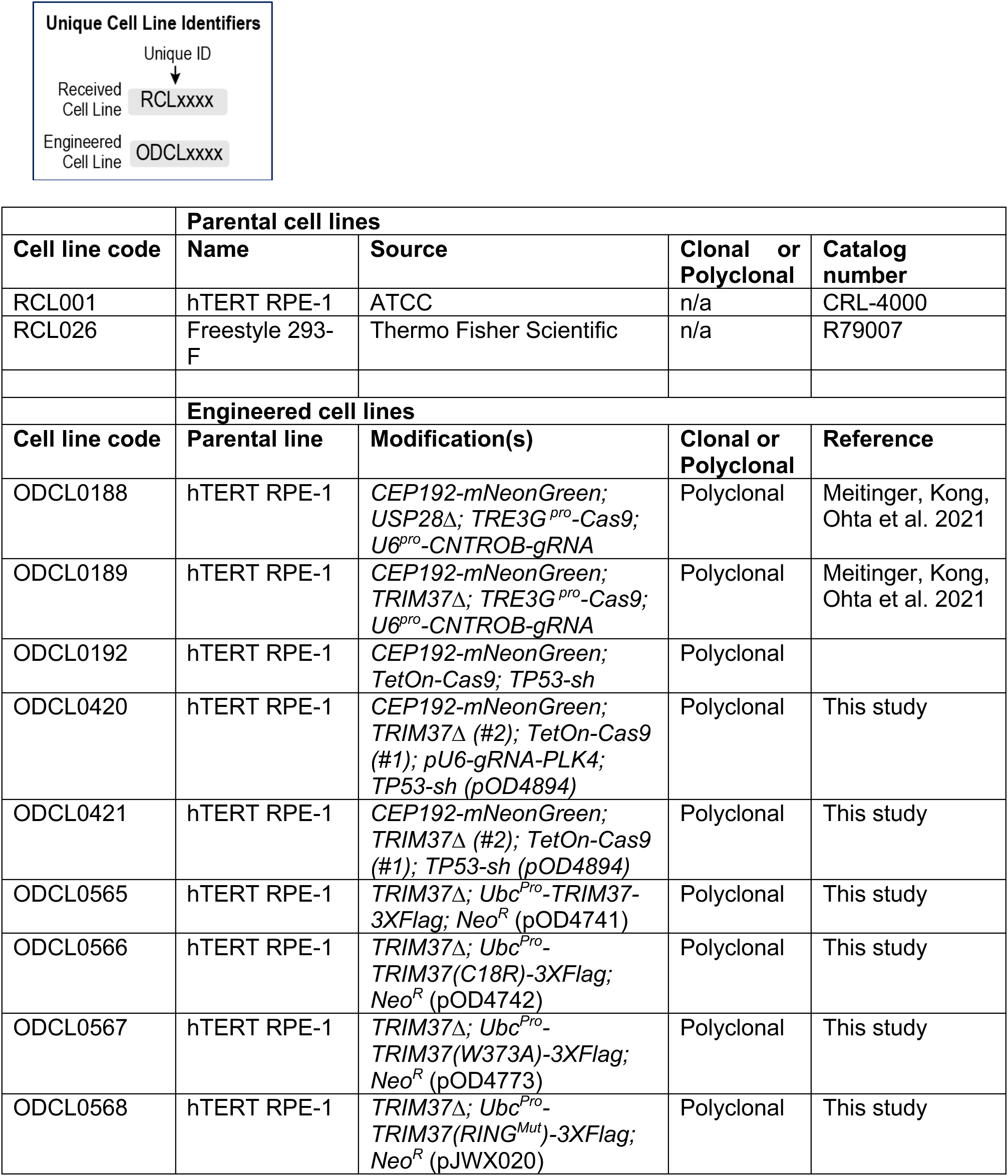
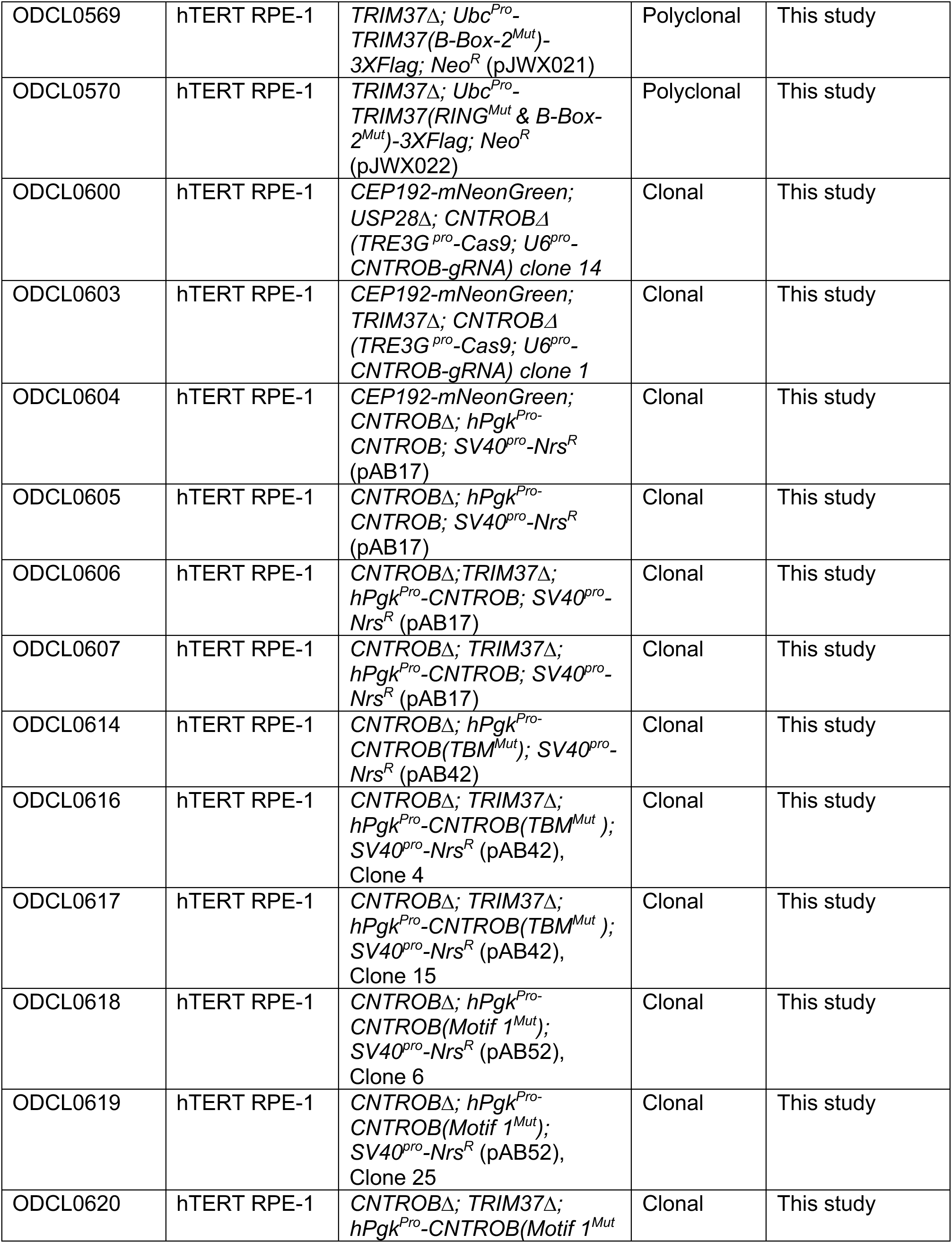
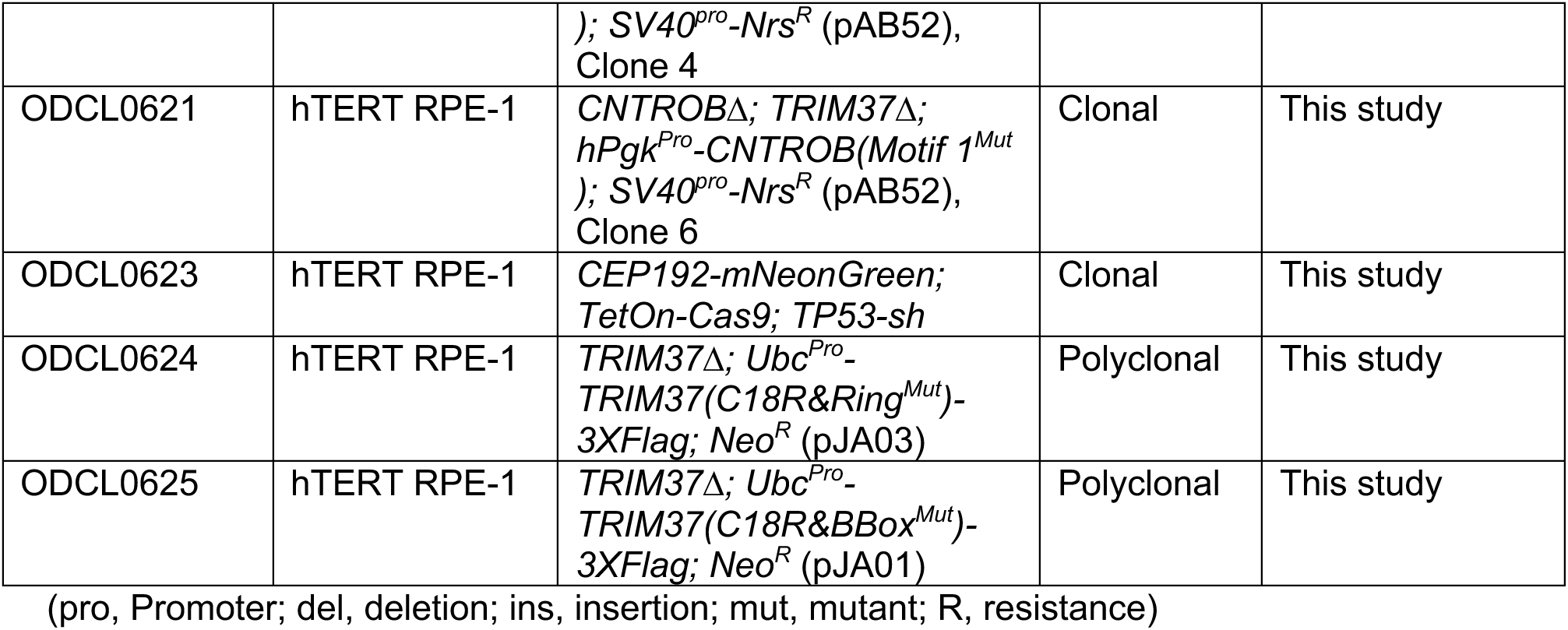
Human cell lines used in this study.

**Table S2.** Plasmids used in this study.

| Plasmid code | Description | Purpose | Bacterial selection | Reference |
| --- | --- | --- | --- | --- |
| pOD3790 | *CMV<sup>pro</sup>-TRIM37-3xFLAG* | Freestyle Expression | Ampicillin | Meitinger et al. 2020 |
| pOD3791 | *CMV<sup>pro</sup>-TRIM37(C18R)-3xFLAG* | Freestyle Expression | Ampicillin | Meitinger et al. 2020 |
| pOD3792 | *CMV<sup>pro</sup>-TRIM37(C18R & W373A)-3xFLAG* | Freestyle Expression | Ampicillin | Meitinger et al. 2020 |
| pOD3797 | *LV-Ubc<sup>pro</sup>-TRIM37-3xFLAG-SV40<sup>pro</sup>-Neo<sup>R</sup>* | Lentiviral Integration | Ampicillin | Meitinger et al. 2020 |
| pOD3798 | *LV-Ubc<sup>pro</sup>-TRIM37(C18R)-3xFLAG-SV40<sup>pro</sup>-Neo<sup>R</sup>* | Lentiviral Integration | Ampicillin | Meitinger et al. 2020 |
| pOD3937 | *LV-U6<sup>pro</sup>-CNTROB-gRNA (lentiGuide-Puro)* | Lentiviral Integration | Ampicillin | Meitinger, Kong, Ohta et al. 2021 |
| pOD3938 | *CMV<sup>pro</sup>-5xMYC-CNTROB* | Freestyle Expression | Ampicillin | Meitinger, Kong, Ohta et al. 2021 |
| pOD4773 | *LV-Ubc<sup>pro</sup>-TRIM37(W373A)-3xFLAG-SV40<sup>pro</sup>-Neo<sup>R</sup>* | Lentiviral Integration | Ampicillin | This study |
| pOD4783 | *CMV<sup>pro</sup>-TRIM37(W373A)-3xFLAG* | Freestyle Expression | Ampicillin | This study |
| pOD4894 | *LV-pLKO1-sh-TP53* | Lentiviral Integration | Ampicillin | Addgene # 19119 |
| pOD4913 | *LV-Addgene#52963 with TRIM37-gRNA* | Lentiviral Integration | Ampicillin | This study |
| pOD5400 | *LIC-miniTRIM37 RBhcc<sup>WT</sup>-StreptII-6xHis* | Bacterial Expression | Ampicillin | This study |
| pOD5401 | *LIC-miniTRIM37 RBhcc<sup>Ringmut</sup>-StreptII-6xHis* | Bacterial Expression | Ampicillin | This study |
| pOD5402 | *LIC-miniTRIM37 RBhcc<sup>B-box-2mut</sup>-StreptII-6xHis* | Bacterial Expression | Ampicillin | This study |
| pOD5403 | *LIC-miniTRIM37 Bhcc<sup>WT</sup>-StreptII-6xHis* | Bacterial Expression | Ampicillin | This study |
| pOD5404 | *LIC-miniTRIM37 Bhcc<sup>B-box-2mut</sup>-StreptII-6xHis* | Bacterial Expression | Ampicillin | This study |
| pOD5405 | *6xHis-MBP-TRIM37 TRAF(274-407)* | Bacterial Expression | Ampicillin | This study |
| pOD5406 | *6xHis-MBP-TRAF<sup>mut</sup>(274-407, W373A)* | Bacterial Expression | Ampicillin | This study |
| pAB001 | *CMV<sup>pro</sup>-5xMYC-CNTROB(1-475)* | Freestyle Expression | Ampicillin | This study |
| pAB003 | *CMV<sup>pro</sup>-5xMYC-CNTROB(460-903)* | Freestyle Expression | Ampicillin | This study |
| pAB005 | *CMV<sup>pro</sup>-5xMYC-CNTROB(1-576)* | Freestyle Expression | Ampicillin | This study |
| pAB007 | *CMV<sup>pro</sup>-5xMYC-CNTROB(1-767)* | Freestyle Expression | Ampicillin | This study |
| pAB017 | *LV-hPgk-mRuby-CNTROB- SV40<sup>pro</sup> Nrs<sup>R</sup>* | Lentiviral Integration | Ampicillin | This study |
| pAB022 | *CMV<sup>pro</sup>-5xMYC-CNTROB(Motif 1<sup>mut</sup>)* | Freestyle Expression | Ampicillin | This study |
| pAB025 | *CMV<sup>pro</sup>-5xMYC-CNTROB(TBM<sup>mut</sup>)* | Freestyle Expression | Ampicillin | This study |
| pAB036 | *CMV<sup>pro</sup>-5xMYC-CNTROB(1-767, TBM<sup>mut</sup>)* | Freestyle Expression | Ampicillin | This study |
| pAB042 | *LV-hPgk-mRuby-CNTROB(TBM<sup>mut</sup>)-SV40<sup>pro</sup>-Nrs<sup>R</sup>* | Lentiviral Integration | Ampicillin | This study |
| pAB052 | *LV-hPgk-mRuby-CNTROB(Motif 1<sup>mut</sup>)-SV40<sup>pro</sup>-Nrs<sup>R</sup>* | Lentiviral Integration | Ampicillin | This study |
| pAB053 | *CMV<sup>pro</sup>-TRIM37(RING<sup>mut</sup>)-3xFLAG* | Freestyle Expression | Ampicillin | This study |
| pAB054 | *CMV<sup>pro</sup>-TRIM37(B-box-2<sup>mut</sup>)-3xFLAG* | Freestyle Expression | Ampicillin | This study |
| pAB055 | *CMV<sup>pro</sup>-TRIM37(RING<sup>mut</sup> & B-box-2<sup>mut</sup>)-3xFLAG* | Freestyle Expression | Ampicillin | This study |
| pAB056 | *CMV<sup>pro</sup>-TRIM37(C18R & RING<sup>mut</sup>)-3xFLAG* | Freestyle Expression | Ampicillin | This study |
| pAB057 | *CMV<sup>pro</sup>-TRIM37(C18R & B-box-2<sup>mut</sup>)-3xFLAG* | Freestyle Expression | Ampicillin | This study |
| pJA001 | *LV-Ubc<sup>pro</sup>-TRIM37(C18R & B-box-2<sup>mut</sup>)-3xFLAG* | Lentiviral Integration | Ampicillin | This study |
| pJA003 | *LV-Ubc<sup>pro</sup>-TRIM37(C18R-RING<sup>mut</sup>)-3xFLAG* | Lentiviral Integration | Ampicillin | This study |
| pJWX020 | *LV-Ubc<sup>Pro</sup>-TRIM37(Ring<sup>Mut</sup>)-3XFLAG-Neo<sup>R</sup>* | Lentiviral Integration | Ampicillin | This study |
| pJWX021 | *LV-Ubc<sup>Pro</sup>-TRIM37(B-Box-2<sup>Mut</sup>)-3XFLAG-Neo<sup>R</sup>* | Lentiviral Integration | Ampicillin | This study |
| pJWX022 | *LV-Ubc<sup>Pro</sup>-TRIM37(RING<sup>Mut</sup>&B-Box-2<sup>Mut</sup>)-3XFLAG-Neo<sup>R</sup>* | Lentiviral Integration | Ampicillin | This study |

(pro, Promoter; LV, lentiviral; LIC, Ligation-independent cloning)

**Table S3.**
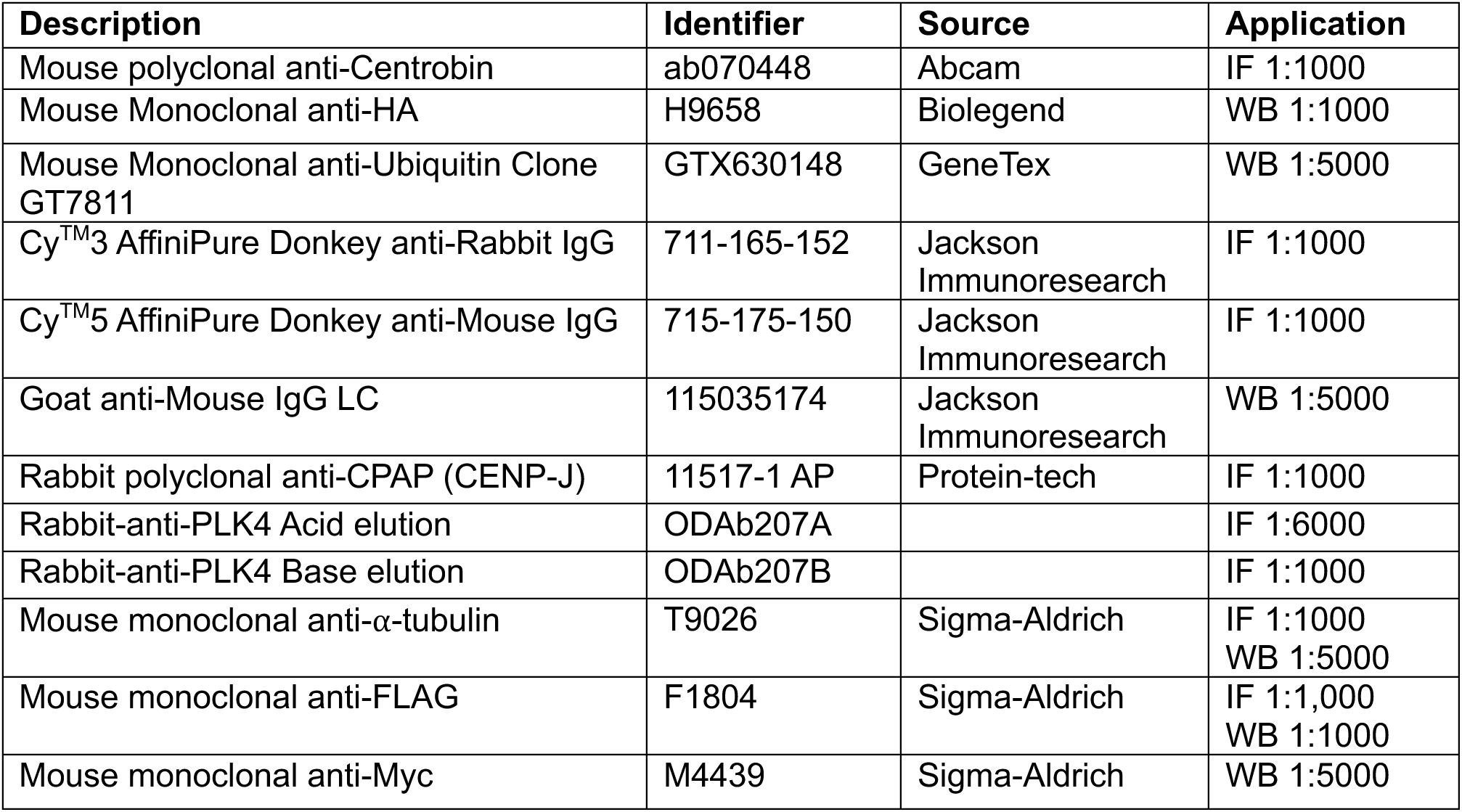
Antibodies used in this study.

